# U-statistics-based Pairwise Discordance Regression for Joint Mean-Variability Inference in Longitudinal Studies with Missing Data

**DOI:** 10.64898/2026.11.04.764284

**Authors:** Jinyuan Liu, Amir Javid, Sarah Osmundson, Xin M. Tu, Jonathan S. Schildcrout

## Abstract

Longitudinal studies are central to scientific inquiry across disciplines; yet, statistical analyses have predominantly focused on mean trajectories while treating variability as a secondary or nuisance component. Increasing evidence suggests that temporal variability and heterogeneity themselves encode critical scientific information, particularly in studies with repeated measurements, heterogeneous populations, or rich covariate structures. The classical mixed-effects and location-scale model family represents variability through latent variable components but relies on strong parametric assumptions and often requires numerical approximation or computationally intensive estimation. Here we introduce the semiparametric pairwise discordance regression, a unified framework for longitudinal data that directly models pairwise contrasts to simultaneously characterize mean trajectories, between-subject heterogeneity, and within-subject temporal variability. By targeting pairwise discordances rather than latent variance components, the proposed approach enables robust inference without specifying full parametric distributions for latent components or error terms, provided the specified conditional moment restrictions hold. Pair-level covariate encodings further accommodate flexible covariate structures and allow variability parameters to be interpreted through pairwise contrasts or dissimilarities. Estimation and inference are carried out via (weighted) U-statistic-based generalized estimating equations (UGEE) with a valid sandwich variance estimator, and we extend the fitting procedure to accommodate missing-at-random dropout. We establish the asymptotic distributions of the resulting estimators under mild regularity conditions. Simulation studies demonstrate reliable finite-sample performance in the settings considered, including non-Gaussian errors and misspecified working correlation structures, whereas likelihood-based parametric fits can be sensitive to distributional assumptions or numerically unstable. An application to longitudinal blood pressure during pregnancy illustrates the framework on real data.

## 1 INTRODUCTION

### 1.1 Background

Classical longitudinal analysis has been dominated by inference on mean trajectories, with variability typically treated as a nuisance component or modeled to improve the efficiency of the estimated mean parameters ^1,2,3^. Increasingly, however, both theoretical and empirical work have demonstrated that variability itself encodes distinct scientific information, reflecting latent heterogeneity, temporal instability, and differential response dynamics that are not revealed from mean trajectories alone ^4,5,6,7^.

This perspective is further underscored by modern longitudinal designs, which are characterized by denser repeated measurements ^8^, greater heterogeneity across individuals ^9,10,11^, and increasingly rich covariate information collected alongside the primary outcome ^12,13^. Together, these factors give rise to complex dependence structures that challenge traditional variance modeling assumptions ^14,15^. More broadly, variability and heterogeneity have emerged as primary inferential targets, rather than as secondary by-products of mean modeling alone ^16,17,18,19,20^.

A substantial body of methodological work has responded to this shift by extending classical mixed-effects models to accommodate heteroscedasticity and covariate-dependent variability. Early contributions relaxed the homogeneous within-subject variance assumptions by linking variance components to covariates through parametric functions, often using quasi-likelihood or method-of-moments approaches to avoid numerical integration ^21,22^. The family of mixed-effects location-scale models (LSMs) further enriched this framework by allowing covariates to influence both mean trajectories (location) and residual variability (scale), yielding a unified parametric approach to characterizing within- and between-subject variability over time ^4,15,23,24,25,26^. These models have been shown to be useful across a wide range of applications, yielding reduced bias and more nuanced inference on dynamic processes, particularly when outcomes exhibit strong temporal instability such as mood, blood pressure, or blood glucose in patients with diabetes ^27,28,29^.

Despite these strengths, classical mixed-effects and location-scale models impose several constraints that are increasingly challenged by modern longitudinal studies. First, heterogeneity is represented through latent subject-level variance components, which impose additive constraints on pairwise heterogeneity that the data need not satisfy. Second, likelihood-based inference typically must specify distributional models for the responses and random effects. Third, although robust or method-of-moments alternatives have been proposed ^25^, these approaches often do not simultaneously address missing data in repeated measures ^30,31^. Finally, likelihood-based estimation for flexible location-scale models can be computationally demanding due to repeated numerical integration or Markov chain Monte Carlo (MCMC), limiting scalability in larger longitudinal settings ^26^. To address these limitations, we shift the target of modeling from latent variance components to observed contrasts. Rather than representing variance through latent random effects, we construct pairwise discordances: differences between outcomes across subjects, across time points, or both. Although defined without reference to any latent structure, these discordances encode between-subject (BS) heterogeneity and within-subject (WS) temporal variability.

To see this, consider a random-intercept linear mixed-effects model ^31^

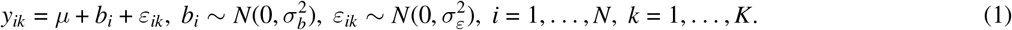

where *y*_*ik*_ is a continuous response of subject *i* at time *k, µ* is the population mean, *b*_*i*_ a subject-specific random effect, and *ε*_*ik*_ the within-subject error. The parameters 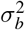 and 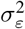 represent between- and within-subject variability, respectively.

These latent variance components can be represented as expectations of observable pairwise contrasts. For any two subjects *i*_1_, *i*_2_ *∈* {1, …, *N*} and two visits *k*_1_, *k*_2_ *∈* {1, …, *K*}:

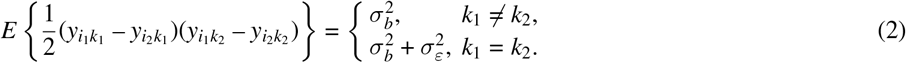

That is, the mean cross-time discordance identifies the between-subject heterogeneity 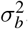, while the mean squared time-specific discordance identifies total variability 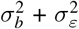. This connection motivates a strategy that treats pairwise discordance as the natural unit of analysis for variability.

In this paper, we propose a pairwise discordance regression framework for longitudinal data that models observable contrasts directly to characterize mean structure, between-subject heterogeneity, and within-subject variability, together with how each component depends on covariates.

### 1.2 Contributions and Organization

This paper contributes to the literature in several ways. First, we introduce a pairwise discordance regression framework for joint mean-variability inference in longitudinal studies. Rather than treating variability as a nuisance component or defining it through latent subject-level variance components, the proposed framework targets observable pairwise functionals that simultaneously characterize marginal mean, between-subject heterogeneity, and within-subject variability.

Second, we establish a formal connection between classical mixed-effects location-scale models and the proposed pairwise discordance framework. We show that a classical location-scale model implies a specific additive structure for pairwise discordance, with between-subject heterogeneity determined by the two subjects’ variance components. Thus, the classical LSM induces a restricted parameterization within the broader pairwise framework. This connection also clarifies two modeling strategies: an *LSM-induced parameterization*, when variability is naturally modeled as an individual characteristic, and a *pair-level parameterization*, when discordance is modeled directly as a function of pair characteristics, including covariate differences between subjects.

Third, we develop U-statistic-based generalized estimating equations (UGEE) for estimation and inference under the resulting model. Because the pairwise system involves no latent variables, estimation requires neither numerical integration nor MCMC. The estimating equations account for dependence induced by overlapping subject pairs, yield sandwich variance estimators based on Hájek projection, and attain the semiparametric efficiency bound within the specified moment-restriction model when the working covariance is correctly specified.

Fourth, we extend the framework to longitudinal missingness through inverse-probability weighted UGEE under monotone missing-at-random (MAR), establishing asymptotic properties for both the complete-data and weighted estimators. Simulation studies evaluate finite-sample calibration, efficiency, and robustness, and benchmark run-time against likelihood-based location-scale models. We provide an `Rcpp` implementation of the proposed estimators, along with a `Python` implementation of the UGEE solver, with open-source code available on `GitHub` (https://github.com/Jinyuanliu00vumc/UGEE-location-scale).

The remainder of the paper is organized as follows. Section 2 formalizes the inferential targets and introduces pairwise functionals through observable discordances. Section 3 develops the pairwise discordance regression framework under pair-level and classical parameterizations and relates it to classical mixed-effects location-scale models, highlighting the structural constraints imposed by latent variance modeling. Section 4 presents estimation and semiparametric inference, including asymptotic linearity, sandwich variance estimation, and extensions to missing data. Sections 5 and 6 evaluate the proposed framework through simulation studies and a real-world application, respectively. Section 7 discusses implications, limitations, and future directions.

## 2 STATISTICAL TARGETS AND PAIRWISE FUNCTIONALS

### 2.1 Notation and Targets of Inference

We begin by formalizing the targets of pairwise modeling and inference. Consider a longitudinal study with *N* subjects, where subject *i* is observed at occasions *k* = 1, …, *K*, common to all subjects, and let *y*_*ik*_ denote a continuous outcome. Longitudinal outcomes may vary both across subjects and within subjects over time ^4^. We therefore distinguish covariates according to the source of variation they are intended to characterize and consider three sets of parameters:

i. Mean-model covariates **x**_*ik*_ characterize the expected outcome trajectory, with corresponding parameters denoted by ***β***.
ii. Subject-level covariates **u**_*i*_ characterize between-subject (BS) heterogeneity, that is, systematic differences in outcome trajectories across individuals. Because BS heterogeneity reflects relatively persistent between-person differences, these effects may admit a trait-like interpretation when supported by the covariates and study design ^32^. Accordingly, **u**_*i*_ typically comprises time-invariant, or time-fixed, covariates, such as sex, treatment group, and baseline measurements, as well as subject-level summaries of time-varying covariates, such as person-specific means. The corresponding parameters are denoted by ***α***.
iii. Time-varying covariates **w**_*ik*_ characterize within-subject (WS) or intra-individual variability, reflecting fluctuations around an individual’s expected trajectory ^9,10^. Their association with WS variability is indexed by ***τ***.

The covariate sets **u**_*i*_ and **w**_*ik*_ may overlap with the mean-model covariates **x**_*ik*_ but need not coincide with them. As in joint mean-variability models ^33,34,4^, ***β, α***, and ***τ*** target the mean trajectory, BS heterogeneity, and WS variability, respectively. In the proposed framework, however, these targets are defined directly through pairwise functionals of the observed outcomes, introduced next.

### 2.2 Pairwise Functionals for Mean and Variability

#### 2.2.1 Mean Functional

Let 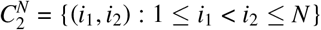 denote the set of unordered subject pairs. For a pair denoted as 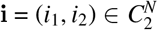, we define the pairwise average outcome at time *k* as

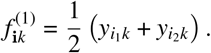

Under a standard mean model involving covariates with 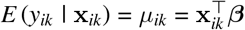, it follows immediately that

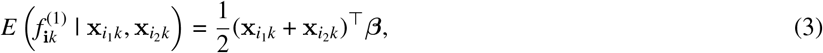

which identifies ***β*** using only observable pairwise quantities.

#### 2.2.2 Lag-1 Discordance Functional

Let 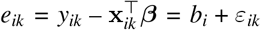 denote the population residuals ^1^. Between-subject heterogeneity reflects persistent differences among individuals, and under the random-intercept specification in (2), it is captured by cross-products of pairwise residual differences. For *k*_1_ *≠ k*_2_,

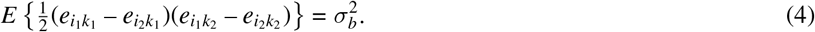

To reduce computational complexity from *O*(*K*^2^) to *O*(*K*), we restrict attention to consecutive occasions and define the lag-1 discordance functional for subject pairs

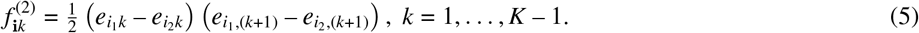

Intuitively, 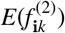 measures the persistence of pairwise residual differences across adjacent visits. A large positive value indicates that subjects with separated residuals at visit *k* tend to have separated residuals in the same relative direction at visit (*k* + 1), reflecting stable between-subject heterogeneity.

This functional generalizes to a lag-*l* discordance 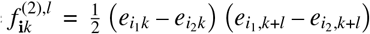, for *l ∈* {1, …, *K* – 1}. Under the random-intercept specification with temporally uncorrelated within-subject errors, these lagged functionals target the same BS variance component, 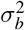, as (4). Restricting attention to lag-1 comparisons can therefore be viewed as an incomplete U-statistic construction ^35^. We use *l* = 1 throughout, as it provides the largest number of comparisons and is straightforward to implement.

#### 2.2.3 Squared Discordance Functional

Under the same specification in (2), the squared residual difference at a single occasion has expectation

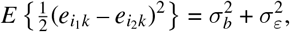

combining both between- and within-subject variance sources. This motivates defining the squared discordance functional

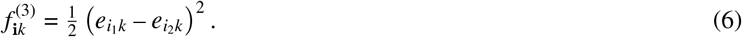

Intuitively, 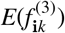 measures how far apart two randomly selected subjects are at a given visit on average; it captures total variability, combining both persistent between-subject differences and transient within-subject fluctuations.

## 3 THE PAIRWISE DISCORDANCE REGRESSION FRAMEWORK

Using the three functionals defined in Section 2.2, we model their conditional expectations through a regression system posed at the pairwise level, treating subject pairs 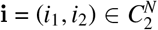 as the units of analysis. We posit the moment restricted semiparametric models ^36^:

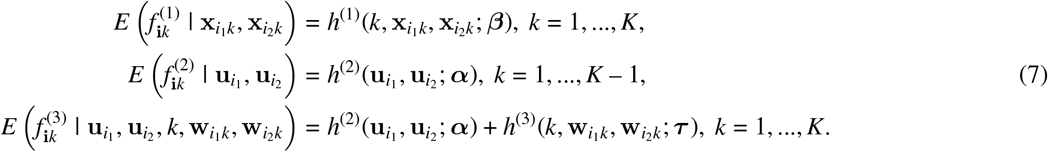

Here, *h*^(1)^(*·*) is usually a smooth function involving time and the pair’s covariates 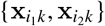, with continuous second-order derivatives ^37^, while *h*^(2)^(*·*) and *h*^(3)^(*·*) may take exponential or other positive-valued links to enforce positivity of the variability components. Throughout, we use the term *discordance* broadly for any such contrast, with the precise form determined by the target of inference.

The system (7) decomposes mean and variability through pairwise functionals, without latent random effects or full distributional assumptions, and collects the parameters of interest in ***θ*** = (***β***^*⊤*^, ***α***^*⊤*^, ***τ*** ^*⊤*^)^*⊤*^. Its flexibility lies in leaving the covariate specifications of *h*^(2)^(*·*) and *h*^(3)^(*·*) open, so each variability component may adopt the parameterization suited to its scientific question. In Section 3.1, we show that the pair-level parameterization directly encodes each pair through functions of the two subjects’ covariates, accommodating categorical and continuous predictors. In Section 3.2, we show that the LSM-induced parameterization combines individual covariate contributions additively across the pair. The two can also be combined, as the real-data application (Section 6) illustrates, yielding both pair-level and subject-level interpretations.

### 3.1 Pair-Level Parameterization

We begin with covariates that are constant within subjects, including intrinsically time-invariant covariates and person-level summaries of time-varying covariates, and then turn to time-varying covariates.

#### 3.1.1 Time-Invariant Covariates and Between-Subject Heterogeneity

Time-invariant covariates **u**_*i*_ are meant to explain persistent subject-level heterogeneity, or between-subject (BS) variability. Their pair-level encodings define the component 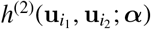 of the discordance regression system (7).

##### 3.1.1.1 Categorical: One-hot Pair Encoding

Consider a binary covariate *u*_*i*_ (e.g., female vs. male). Its full pair-level encoding has three levels, (*ϕ*_*FF*_, *ϕ*_*MM*_, *ϕ*_mix_)^*⊤*^, where *ϕ*_*FF*_ = 1, *ϕ*_*MM*_ = 1, and *ϕ*_mix_ = 1 indicate female-female, male-male, and mixed-sex pairs, respectively. In regression models, taking the female-female pair as the reference level, we use the reduced encoding ***ϕ***(**u**_**i**_) = (*ϕ*_*MM*_, *ϕ*_mix_)^*⊤*^ and specify

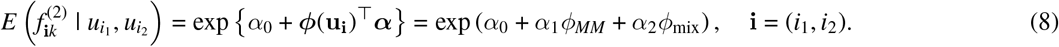

Here exp(*α*_1_) and exp(*α*_2_) represent between-subject heterogeneity ratios for male-male and mixed-sex pairs, respectively, relative to the female-female reference.

More generally, for a categorical covariate with *L* levels, let *U*_*L*_ = {(*l*_1_, *l*_2_) : 0 *≤ l*_1_ *≤ l*_2_ *≤ L* – 1} index the unordered pairings of levels, so that 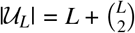. Writing **u**_**i**_ = (*u*_*i*_, *u*_*i*_) for the covariate values of the pair **i** = (*i*_1_, *i*_2_), the pair-level covariate is represented by the one-hot encoding ^38^

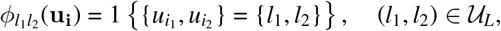

corresponding to a pair at homogeneous levels (*l*_1_ = *l*_2_) or heterogeneous levels (*l*_1_ < *l*_2_). For the latter, this indicator equals one for either ordering, namely 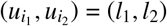 or (*l*_2_, *l*_1_).

For regression, one pairing is chosen as the reference and absorbed into the intercept. Taking (0, 0) pairing as the reference without loss of generality, and ordering the remaining pairings in (*l*_1_, *l*_2_), we define a reduced encoding vector

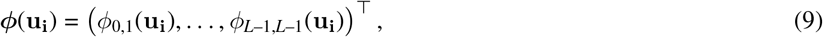

which contains one indicator for each remaining unordered pairing and has length 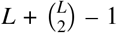. This vector can then enter the conditional mean model for the pairwise discordance functional through 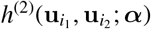 When scientifically appropriate, subject-level categories may be merged *a priori* into coarser levels before pair encoding, to reduce the dimension of ***ϕ***(**u**_**i**_) and enhance the interpretability ^37^.

##### 3.1.1.2 Continuous: Dissimilarity Mapping

When **u**_*i*_ contains continuous or high-dimensional covariates, between-subject heterogeneity may be modeled through pairwise dissimilarities of these covariates ^39^. Such mappings can be constructed at different levels of aggregation. For each covariate or a prespecified set of related covariates, separate pairwise mapping functions can be defined. Let **u**_*ir*_ denote the *r*th covariate or covariate set, we define

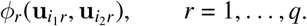

Then, *ϕ*_*r*_(*·, ·*) may be an absolute difference for a scalar covariate or a multivariate dissimilarity for a covariate set. These mappings enter the between-subject model:

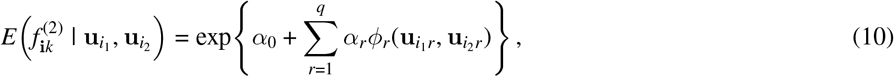

which preserves covariate- or covariate-set-specific interpretations for between-subject heterogeneity.

Alternatively, when the covariate features are naturally viewed jointly, as with neuroimaging, genomic, or sensor-derived measurements, the entire pair 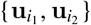 can be summarized by a single dissimilarity functional ^40,37^. For example, using the Euclidean distance

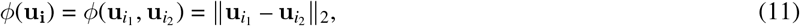

gives

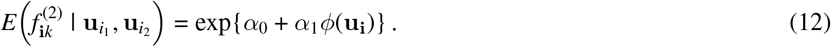

This aggregate representation can be applied directly to high-dimensional covariates without requiring prior feature selection or dimension reduction, while inference continues to target finite-dimensional regression parameters. The choice of *ϕ*(*·, ·*) should therefore reflect both variable type and scientific context: Euclidean or Mahalanobis distances capture global multivariate differences, whereas feature-weighted distances can emphasize selected variables ^41^.

In summary, time-invariant covariates are handled by mapping categorical variables through the one-hot pair encoding in (9) and continuous variables through the pairwise dissimilarity metrics. Combining these categorical and continuous mappings gives the pair-level covariate vector ***ϕ***(**u**_**i**_), which specializes *h*^(2)^(*·*) in (7) to

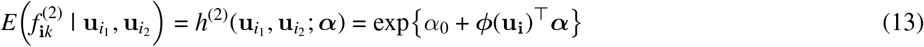

and supplies the between-subject (BS) terms in the discordance regression.

#### 3.1.2 Time-Varying Covariates and Total Variability

Time-varying covariates for each subject **w**_*ik*_ fluctuate over time *k* = 1, …, *K*. Their pair-level construction parallels Section 3.1.1, applied separately at each occasion: categorical components are mapped through the pair encoding in (9) evaluated at 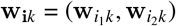, and continuous components through a pre-specified distance functional 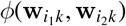.

Combining them gives the pair-level covariate vector ***ϕ***(**w**_**i***k*_), which, unlike its time-invariant counterpart, varies with *k*. These within-subject components, together with time, specify *h*^(3)^(*·*) in (7) as

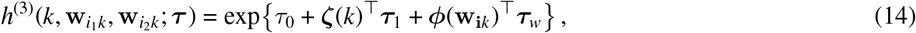

where ***ζ***(*·*) is a pre-specified function of time (e.g., linear, piecewise, or indicator). Under this parameterization, exp(*τ*_0_) is the within-subject variability at the reference time and reference covariate values, ***τ***_1_ captures its temporal trend, and ***τ***_*w*_ quantifies how pairwise dissimilarity in time-varying covariates contributes to within-subject temporal variability.

The within-subject variability component *h*^(3)^(*·*) and the between-subject component *h*^(2)^(*·*) together define the total variability through the conditional mean of the squared discordance functional in (6):

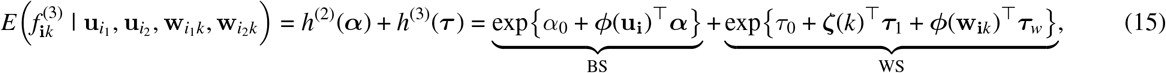

where the exponential links keep both components positive.

This decomposition reflects the scientific distinction between two sources of longitudinal variation. Between-subject heterogeneity represents stable individual differences that persist across occasions, whereas within-subject variability may change over time or with changes in covariates. Accordingly, the within-subject component varies across time through ***ζ***(*k*) and with pairwise covariate dissimilarity through ***ϕ***(**w**_**i***k*_), while the between-subject component is specified as the same *h*^(2)^(*·*) term in (13), does not depend explicitly on *k*, and shares ***α*** across the lag-1 and squared-discordance moment equations, so the between-subject term in (15) is identified by the lag-1 moments rather than estimated freely from the squared discordances.

##### Remark 1.

*Parameter Interpretation*. Taken together, the pair-level parameterizations shift the inferential target from individuals to subject pairs. For time-invariant covariates, under the categorical encoding in (9), the parameter 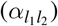 is the ratio of expected between-subject variability for a pair of type (*l*_1_, *l*_2_) to that for the reference pairing. For continuous covariates represented through the dissimilarity mapping in (10) or (12), a one-unit increase in the covariate dissimilarity multiplies the expected between-subject variability by exp(*α*_*r*_); thus, *α*_*r*_ > 0 indicates greater outcome heterogeneity among more covariate-dissimilar pairs.

Analogously, time-varying covariates enter the within-subject variability component through pair-level encodings that vary by occasion. For a categorical time-varying covariate, 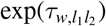 is the ratio of expected within-subject variability for pair type (*l*_1_, *l*_2_) relative to the reference pairing, holding time and other covariates fixed. In a crossover design with time-varying treatment status, for example, if control-control pairs are the reference, then exp(*τ*_*w*,*TT*_) quantifies treatment-associated stabilization or destabilization of within-subject trajectories by comparing occasions when both members of the pair are under treatment with occasions when both are under control. For continuous time-varying covariates, each component of ***τ***_*w*_ gives the multiplicative change in expected within-subject variability for a one-unit increase in pairwise covariate dissimilarity, while ***τ***_1_ describes the temporal trend through ***ζ***(*·*). For example, when ***ζ***(*k*) = *k*, exp(*τ*_1_) is the multiplicative change in within-subject variability per unit increase in *k*.

### 3.2 LSM-Induced Parameterization

Gaussian mixed-effects location-scale models (LSMs) are widely used for joint mean-variance modeling in modern longitudinal data, particularly in heterogeneous populations or studies with rich covariate structures ^4,25,26^. We now show that the pairwise moment structure induced by an LSM is a special case of the proposed discordance regression system in (7). Specifically, the LSM imposes an additive parameterization on the expected discordance functionals through particular choices of *h*^(2)^(*·*) and *h*^(3)^(*·*).

#### 3.2.1 The Gaussian Location-Scale Model

To attribute variability to covariates, a mixed-effects location-scale model extends the random-intercept model in (1) by allowing both between- and within-subject variances to depend on covariates ^4^:

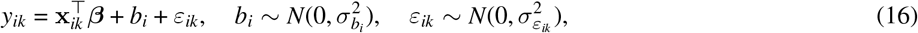

with variance components specified, for example, as

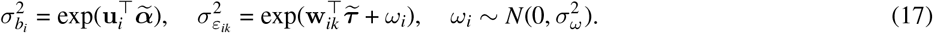

The exponential links ensure positivity of the variance components. Unlike the classical mixed-effects model, (16)-(17) involves two random effects: *b*_*i*_, the random *location* effect governing the individual mean trajectory, and *ω*_*i*_, the random *scale* effect modulating within-subject variability.

In this parameterization, 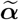 links subject-level covariates **u** to the between-subject variance 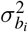 and 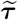 links time-varying covariates **w**_*ik*_ to the within-subject variance 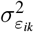. The variance of random scale effect 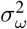 captures residual within-subject variability beyond the observed covariates.

We denote the full parameter vector induced by LSM as 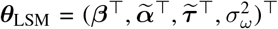. The tilde marks subject-level parameters, which parameterize individual variance components and in turn induce pairwise moments, and distinguishes them from the pair-level ***α*** and ***τ*** of Section 3.1, which model expected discordance directly as a function of pair-level encodings or dissimilarities.

#### 3.2.2 Pairwise Moments Induced by the LSM

Next, we derive the pairwise moment structure induced by the LSM and show which moment targets are identified by the proposed discordance functionals, without requiring distributional assumptions beyond those already imposed by the LSM.

##### 3.2.2.1 Mean Functional

For a subject pair **i** = (*i*_1_, *i*_2_) at occasion *k*, the mean functional satisfies

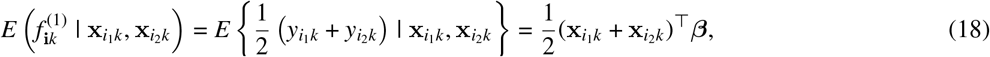

which recovers ***β*** from the mean structure of (16). This identity holds irrespective of the distributional assumptions imposed on *b*_*i*_, *ε*_*ik*_, or *ω*_*i*_, and likelihood-based estimation is therefore not required for identifying this marginal mean parameter ***β***.

##### 3.2.2.2 Between-Subject Heterogeneity

Recall the lag-1 discordance functional 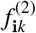 in (5) measures the persistence of pairwise residual differences across adjacent visits and corresponds to the between-subject heterogeneity. Under the covariate-dependent LSM in (17), one can show that

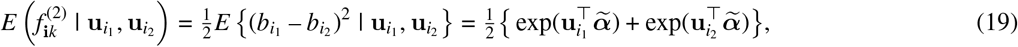

since 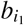 and 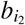 are assumed uncorrelated for *i*_1_ *≠ i*_2_. Thus, 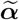 is identified through this moment condition, without specifying full distributional forms for the random effects.

##### 3.2.2.3 Total Variability and Decomposition

Similarly, recall the squared discordance functional 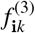 measures the total variability at visit *k*, combining both between-subject heterogeneity and within-subject variability. Under the LSM, its conditional mean gives

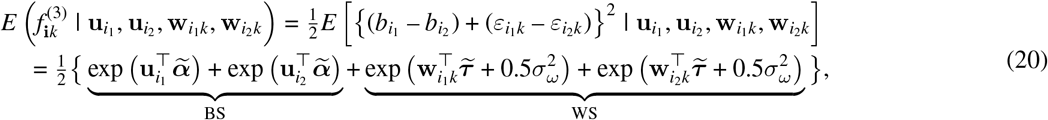

since the within-subject errors are assumed mean zero and uncorrelated across visits and with the random location effects ^2,4^.

The first two terms coincide with the between-subject contribution identified by the lag-1 discordance in (19), whereas the latter two terms capture within-subject variability. The additive term 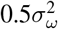 arises from marginalizing over the random scale effect, since 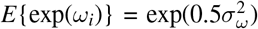 under the LSM assumption that 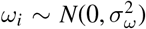. Under this parameterization, the interpretation follows the LSM, where a one-unit increase in the *j*th component of **u**_*i*_ multiplies that subject’s between-subject variance by 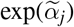, and a one-unit increase in the *j*th component of **w**_*ik*_ multiplies the within-subject variance by 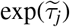, holding other covariates fixed.

### 3.3 Comparing the Two Parameterizations

The LSM arises as a restricted subclass of the proposed pairwise discordance framework. The framework in (7) defines a joint moment system in which the marginal mean determines the first-order component, while *h*^(2)^(*·*) and *h*^(3)^(*·*) characterize between-subject (BS) and within-subject (WS) variability through flexible covariate parameterizations. The LSM moments in (19) and (20) correspond to particular choices of these functions. Thus, recovering the LSM amounts to imposing restrictions on *h*^(2)^(*·*) and *h*^(3)^(*·*).

For between-subject heterogeneity, the LSM specifies the subject-specific variance 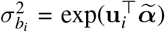. For a pair of subjects, these variance components enter additively as in (19); hence, pairwise heterogeneity depends on the two subjects only through their summed variance contributions. Pair-level specifications such as (11) relax this restriction by modeling discordance directly as a function of pair characteristics.

The restriction is particularly transparent for a binary covariate *u*_*i*_ *∈* {0, 1}. Under the LSM,

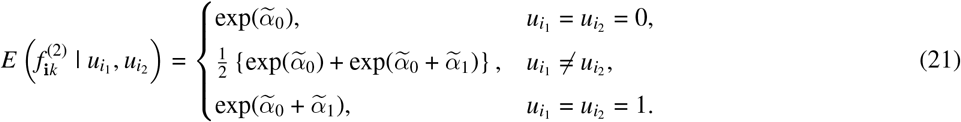

Thus, the heterogeneous cell is constrained to the arithmetic mean of the two homogeneous cells by the additive LSM variance structure.

In contrast, consider the pair-level encoding ***ϕ***(**u**_**i**_) = (*ϕ*_11_, *ϕ*_01_)^*⊤*^, where the subscripts index unordered pairs of covariate levels and *u*_*i*_ = *u*_*i*_ = 0 is the reference cell. The pairwise model specifies 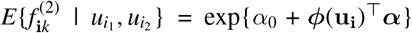. The homogeneous cells reproduce those in (21) under the reparameterization *α*_0_ = *α*_0_ and *α*_1_ = *α*, whereas the heterogeneous cell,

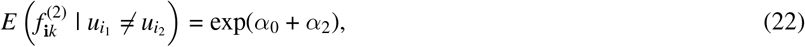

is estimated independently. Equating this with the LSM heterogeneous cell in (21) yields 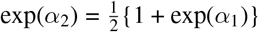. Thus, the LSM restriction is testable within the pair-level parameterization.

#### Remark 2.

*Parameter Dimension*. For a categorical covariate with *L* levels, the LSM imposes 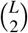 analogous restrictions, one for each heterogeneous pairing, with each heterogeneous cell fixed at the arithmetic mean of its corresponding homogeneous cells. The pair-level parameterization instead estimates these cells directly, allowing the LSM restrictions to be examined from the data.

For continuous covariates, the relative parameter dimension depends on the chosen pair-level representation. A *q*-dimensional subject-level covariate **u**_*i*_ requires *q* coefficients in (17). A pair-level specification may use *q* componentwise dissimilarities or aggregate them through a pairwise dissimilarity measure such as (11). Aggregation can reduce the number of discordance parameters but requires a scientifically meaningful metric ^37^. Thus, neither parameterization is uniformly more parsimonious; the relative dimension depends on the covariate structure and pair-level representation.

#### Remark 3.

*Identification*. For the within-subject (WS) component, the pair-level parameterization directly targets *τ*_0_ in (14), the log WS variability at the reference occasion and covariate values, through the pairwise moment conditions. Under the LSM representation in (20), 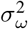 enters through 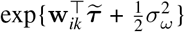. The pairwise moments therefore identify 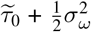, but not 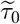 and 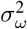 separately. Their separate estimation under the LSM follows from the full parametric model and likelihood, including the assumed distribution of the random scale effect *ω*_*i*_ ^4^. Apart from this decomposition, the corresponding moment targets are identified within the pairwise discordance system without specifying a full outcome distribution.

The two parameterizations also differ in their inferential requirements. Likelihood-based estimation of the LSM proceeds under specified distributional models for the responses and random effects and may require numerical integration or Markov chain Monte Carlo ^4,26^. In contrast, the proposed framework estimates the discordance moments semiparametrically, without specifying a full outcome distribution, and admits computationally simpler estimation as developed in Section 4.

## 4 ESTIMATION AND SEMIPARAMETRIC INFERENCE

Here we develop estimation and inference for the proposed pairwise discordance regression in (7). Two challenges distinguish this setting from classical generalized estimating equations (GEE) ^42^. First, the observational unit is a subject pair; the estimating equations involve sums over pairwise functionals whose summands are dependent due to shared subjects across pairs ^38,43^. Second, because the summands are not independent, classical central limit theorems do not apply directly, and asymptotic normality requires an argument based on Hájek projections of the U-statistic scores onto subject-level contributions ^44,45,46^. To address these challenges, we introduce a U-statistic-based system of generalized estimating equations (UGEE) and derive its asymptotic properties accordingly.

### 4.1 Estimating Equations and U-Statistic Structure

Let 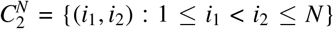 denote the set of unordered subject pairs. For each pair 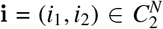, recall that the pairwise functionals 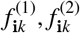, and 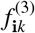 capture the mean, between-subject, and total variability components, respectively. Define the stacked pairwise functional vector and its corresponding conditional mean as

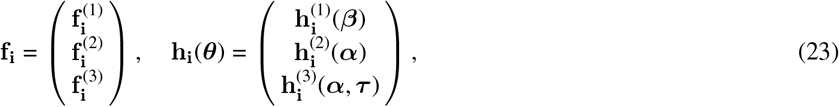

where 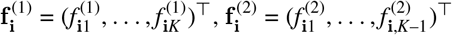, and 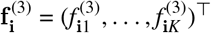.

Let *ℱ*_**i**_ denote the *σ*-algebra generated by all covariates entering the pairwise conditional mean **h**_**i**_(***θ***), including 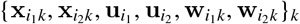. The inferential target ***θ*** = (***β***^*⊤*^, ***α***^*⊤*^, ***τ*** ^*⊤*^)^*⊤*^ is defined as the unique solution to the stacked conditional moment restriction

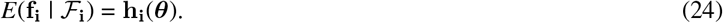

The model is likelihood-free and semiparametric: inference for ***θ*** is based solely on the moment condition (24), with no further assumptions imposed on the joint or marginal distributions of **f**_**i**_.

Define the residual vector **S**_**i**_ = **f**_**i**_ – **h**_**i**_(***θ***) and the Jacobian **G**_**i**_ = (*∂*/*∂****θ***)**h**_**i**_(***θ***)^*⊤*^, of dimension *p ×* dim(**f**_**i**_). We estimate ***θ*** by solving the U-statistic-based generalized estimating equation (UGEE)

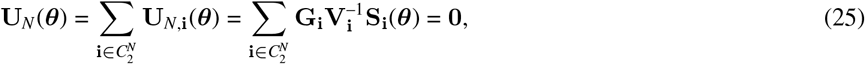

where **V**_**i**_ is a working covariance matrix for **f**_**i**_.

In practice, we adopt a block-diagonal working covariance

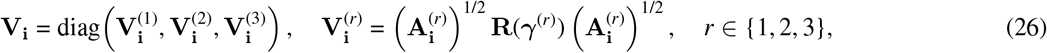

where 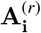 is a diagonal matrix of working marginal variances of 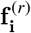, specified as functions of the mean implied by the moment model (24); **R**(***γ***^(*r*)^) is a working correlation matrix across occasions within a pair, of dimension *K* for *r* = 1, 3 and *K* – 1 for *r* = 2; and ***γ*** = (***γ***^(1)*⊤*^, ***γ***^(2)*⊤*^, ***γ***^(3)*⊤*^)^*⊤*^ stacks the working correlation parameters.

For example, under a log-linear model with a Poisson-type variance-mean relationship ^36^, we may take 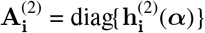. In simulations, **R**(***γ***^(*r*)^) was taken to be independence, exchangeable, AR(1), or the model-based structure implied by a Gaussian location-scale model. For the exchangeable and AR(1) forms, ***γ***^(*r*)^ is estimated by the usual moment estimators applied to the block-*r* Pearson residuals 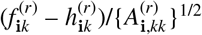, averaged over subject pairs ^47^. Analogous to GEE, consistency of the UGEE estimator 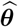 relies only on the moment restriction (24) and does not require correct specification of **V**_**i**_(***γ***).

Because subject pairs may overlap, the summands in (25) are not independent (e.g., pairs **i** = (1, 2) and **i**^*′*^ = (1, 3) are dependent as both involve subject 1), precluding direct application of classical independence-based central limit theorems. Instead, **U**_*N*_ (***θ***) is a multivariate U-statistic whose asymptotic behavior is characterized via its Hájek projection, which we establish next.

### 4.2 Asymptotic Linearity and Sandwich Variance

Let ***θ***_0_ denote the true parameters. Since **G**_**i**_ = (*∂*/*∂****θ***)**h**_**i**_(***θ***) and **V**_**i**_ are *ℱ*_**i**_-measurable, the sensitivity matrix is

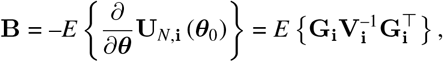

which is assumed nonsingular. Under standard regularity conditions ^46^, including continuous differentiability of **h**_**i**_(***θ***) in a neighborhood of ***θ***_0_ with a dominated Jacobian (Conditions B1–B2) and finite second moments of **S**_**i**_(***θ***_0_) (see **Supplement** for full details), a Taylor expansion of 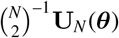 around ***θ***_0_ yields the asymptotic linear (AL) form

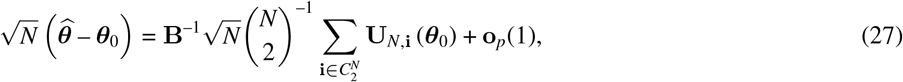

where 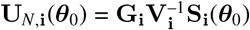. The corresponding influence function (IF) is

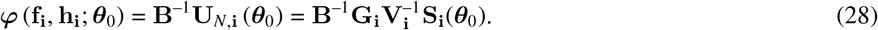

By exploiting equivalence classes in the Hilbert tangent space, it has been established separately that when **V**_**i**_ equals the true conditional covariance of **f**_**i**_ given *ℱ*_**i**_, the influence function in (28) coincides with the efficient influence function (EIF) for ***θ*** under the moment restriction model (24) ^43^, so its variance equals the semiparametric efficiency bound ^36^.

The asymptotic variance of the leading term in (27) is determined by its Hájek projection onto subject-level contributions. For a pair **i** = (*i*_1_, *i*_2_), let 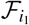 denote the *σ*-algebra generated by subject *i*_1_’s full longitudinal data (outcomes and covariates), and let *I*_2_ *∈* {1, …, *N*} \ {*i*_1_} be an independent subject index. Define the subject-level projected score

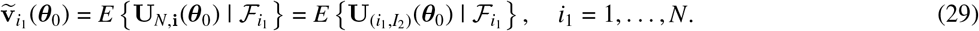

The effective sample size is therefore *N* rather than 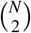, and the estimator achieves the 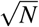 rate, as summarized below.

**Theorem 1** (UGEE asymptotics and sandwich variance). Assume: (i) the moment restriction *E*(**f**_**i**_ | *ℱ*_**i**_) = **h**_**i**_(***θ***) holds under ***θ***_0_; (ii) **h**_**i**_(***θ***) is continuously differentiable in a neighborhood of ***θ***_0_; (iii) 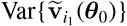 exists and is nonsingular. Let 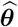 solve the UGEE in (25) using any 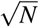-consistent estimator of the working covariance parameter(s). Then

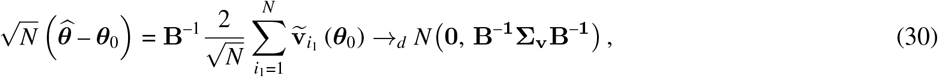

where 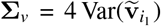. Consistent estimators of **B** and **Σ**_*v*_ are obtained via empirical averages of pairwise scores and their Hájek projections, yielding the sandwich variance estimator 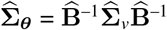.

### 4.3 Missing Data and Inverse Probability Weighting

Missing data arise commonly in longitudinal studies and require careful handling within the pairwise U-statistic framework: when one or both subjects in a pair are unobserved at a given occasion, the corresponding pairwise functional is unobserved, and complete-pair analysis may yield biased inference under missing at random (MAR) ^30^. We extend the UGEE in (25) using inverse probability weighting (IPW) under monotone dropout, a common missingness mechanism in longitudinal settings ^36,48^.

Let *r*_*ik*_ *∈* {0, 1} indicate whether *y*_*ik*_ is observed at time *k*, and let 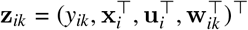 collect all variables relevant for missingness modeling, with **z**_*ik*–_ = {**z**_*i*1_, …, **z**_*i*,(*k*–1)_} denoting the observed history up to time (*k* – 1). We assume: (i) missing at random (MAR) conditional on the observed history; (ii) independence of missing processes across subjects (non-interference); (iii) positivity, so that the probability of being observed at each time point is bounded away from zero; and (iv) monotone dropout, where *r*_*i*,(*k*+1)_ = 1 implies *r*_*ik*_ = 1. Under (i)-(iv), the missing process is modeled via a logistic one-step transition model for subject *i* remaining observable from time (*k* – 1) to *k*:

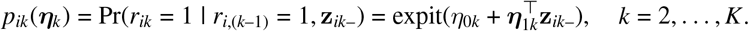

The cumulative retention probability that subject *i* remain observed through time *k* is then

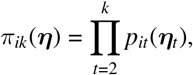

where ***η*** denotes the parameter vector for the missingness model, and *π*_*i*1_ ≡ 1 since all subjects are observed at baseline. Extensions to non-monotone missingness are possible with appropriately constructed missingness models ^36^; however, the implementation and empirical analyses in this paper focus on monotone dropout.

The mean functional 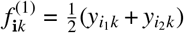 decomposes additively into subject-level contributions, so subject-level IPW applies naturally. We define the weighted mean functional as

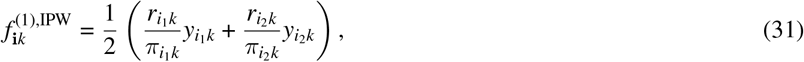

where each subject’s observed outcome is upweighted by the inverse of their retention probability. Under MAR and correct specification of the missingness model,

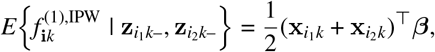

so partially observed pairs still contribute information for estimating ***β***. When one subject is observed and the other is missing, the observed subject’s contribution is upweighted while the missing subject contributes zero; this reduces efficiency relative to complete pairs but preserves unbiasedness under MAR.

The variability functionals 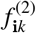 and 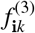 involve products of outcomes across subjects and, for 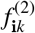, across adjacent time points. Unlike the mean functional, they do not decompose additively into subject-level contributions and cannot be recovered from a single subject’s data alone. Subject-level IPW is therefore insufficient, and weighting must be applied at the pair-level to correct for the selection bias induced by restricting to complete pairs.

Under the non-interference assumption (ii), the joint probability that both subjects in a pair are observed factorizes as

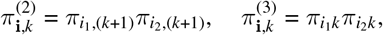

where 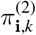 involves retention probabilities at time *k* + 1 rather than *k*, reflecting the fact that the lag-1 functional 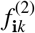 depends on outcomes at both occasions *k* and *k* + 1 as in (5). The pairwise observation indicators are

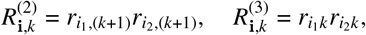

and the corresponding pair-level IPW weights and weighted functionals are

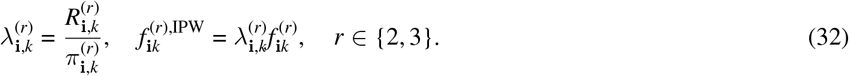

Under MAR and correct model specification,

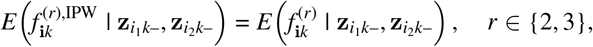

so the weighted variability functionals recover the same conditional expectations as in the complete data. The complete-data UGEE in (25) is extended in Section 4.4 by substituting these IPW functionals and jointly estimating the target parameters ***θ*** and the missingness-model parameters ***η***.

### 4.4 Joint Weighted Estimating Equation

Because the observation probabilities *π* are unknown and must be estimated from data, treating 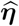 as fixed would ignore the uncertainty introduced by estimating the missingness model and yield an invalid variance estimator for 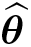^49,50^. Valid inference therefore requires joint estimation of the outcome and missingness parameters, so that the sandwich variance propagates uncertainty from the missingness model to inference for ***θ*** ^51^.

Let 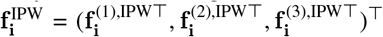 denote the stacked weighted functionals across time points, where subject-level weights are applied to the mean component and pair-level weights to the variability components, as constructed in Section 4.3. The outcome estimating function is obtained by substituting 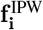 into (25):

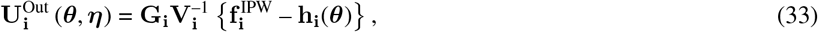

where **G**_**i**_ and **V**_**i**_ are as defined in (25) and (26). Under MAR and correct missingness model specification, this estimating function is unbiased: 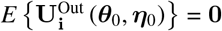.

To construct the missingness estimating equation on the same pairwise index set as (33), we work with the induced pair-level missing process. Let 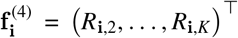 denote the pairwise observation indicators 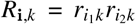, with *R*_**i**,1_ *≡* 1, and define 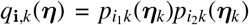 as the induced pair-level transition probabilities, with **q**_**i**_(***η***) =(*q*_**i**,2_, …, *q*_**i**,*K*_)^*⊤*^. Letting **Q**_**i**_ = (*∂*/*∂****η***)**q**_**i**_(***η***)^*⊤*^, the missingness estimating function is

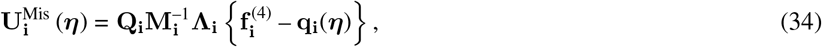

where **M**_**i**_ = (**A**_**i**_)^1/2^**R**(***γ***)(**A**_**i**_)^1/2^ is a working variance, with **A**_**i**_ diagonal with entries *q*_**i**,*k*_(***η***){1 – *q*_**i**,*k*_(***η***)}, conformable with the binary indicators *R*_**i**,*k*_, and **Λ**_**i**_ = diag(*R*_**i**,1_, …, *R*_**i**,*K*–1_) restricts each component to pairs still at risk at the preceding occasion, where the transition model is defined. Under the induced pairwise model, (34) is unbiased: 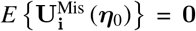, by iterated expectation over the at-risk set.

Stacking the outcome and missingness components yields the joint estimating equation

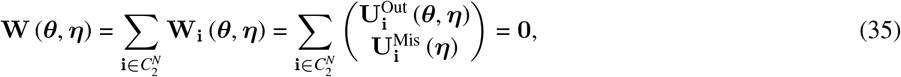

with solution 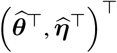.

**Theorem 2** (Joint asymptotics with estimated weights). Under Conditions A-F in the **Supplement** and assuming 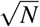-consistent estimation of working correlation parameters, the joint estimators 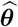 and 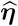 are consistent and asymptotically normal:

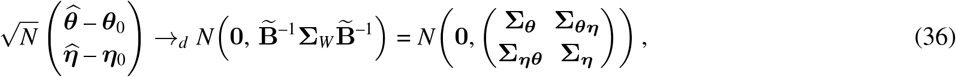

where 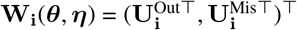 denotes the pair-level contribution to (35), and

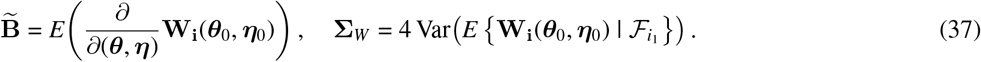

Inference for ***θ*** is based on the upper-left block **Σ**_***θ***_ in (36). Its estimator 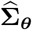, the upper-left block of 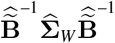, is a consistent estimator of **Σ**_***θ***_. Importantly, **Σ**_***θ***_ differs from the variance obtained by treating 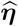 as fixed, as it correctly propagates the uncertainty from estimating the missingness model to inference for ***θ***; full details are given in the **Supplement**.

For testing *H*_0_ : **C*θ***_0_ = **0**, where **C** is a contrast matrix of rank *c*, the Wald statistic

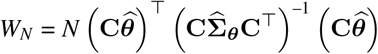

follows an asymptotic 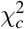 distribution under *H*_0_.

## 5 SIMULATION STUDIES

We conducted simulation studies to evaluate the finite-sample performance of the proposed methods along three dimensions: 1) parameter recovery and inference under correct specification, 2) efficiency relative to likelihood-based inference when the Gaussian model is correctly specified, and 3) robustness when the Gaussian assumptions fail. Two data-generating designs were considered. Design A supported the calibration study in Section 5.2, where we assessed parameter recovery for UGEE and weighted UGEE (WUGEE), as well as the validity of sandwich variance estimation. Design B supported the efficiency study in Section 5.3 and the robustness study in Section 5.4, both of which compare UGEE with the likelihood-based LSM. All settings used 500 Monte Carlo replications and a two-sided significance level of *α* = 0.05. Computations were conducted in `R`, with intensive components implemented in `Rcpp` ^52^.

### 5.1 Simulation Design

We summarize the two simulation designs in **Table 1**. Design A used *K* = 5 repeated measurements and *N* = 150 or 500 for the primary calibration study, with results for *N* = 300 reported in **Supplemental Table S1**. Design B fixed *N* = 300 while varying the number of repeated measurements over *K* = 5 and 10 to evaluate efficiency and robustness.

**TABLE 1.**
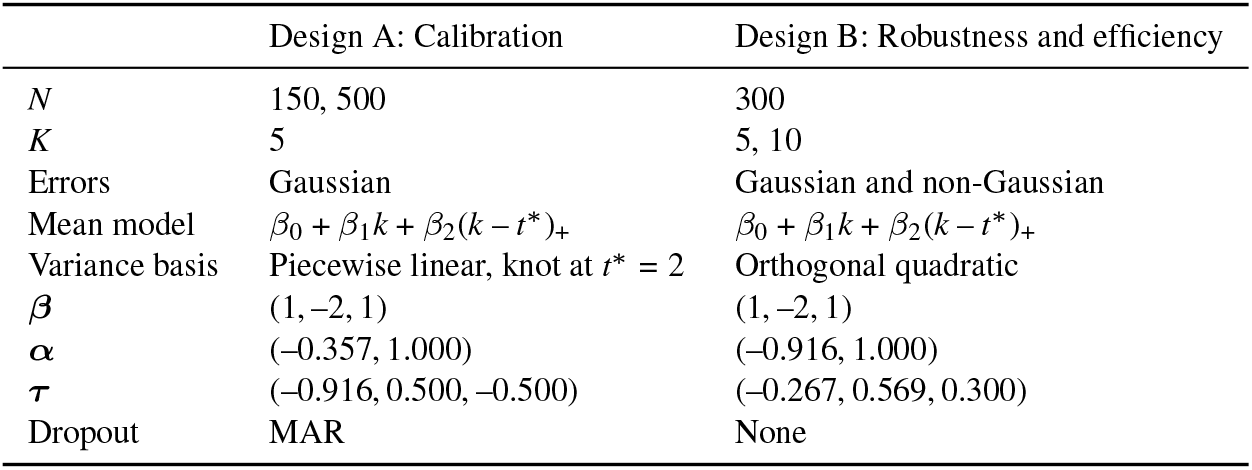
Summary of simulation designs.

|  | Design A: Calibration | Design B: Robustness and efficiency |
|---|---|---|
| *N* | 150, 500 | 300 |
| *K* | 5 | 5, 10 |
| Errors | Gaussian | Gaussian and non-Gaussian |
| Mean model | *β*<sub>0</sub> + *β*<sub>1</sub>*k* + *β*<sub>2</sub>(*k* − *t*<sup>\*</sup>)<sub>+</sub> | *β*<sub>0</sub> + *β*<sub>1</sub>*k* + *β*<sub>2</sub>(*k* − *t*<sup>\*</sup>)<sub>+</sub> |
| Variance basis | Piecewise linear, knot at *t*<sup>\*</sup> = 2 | Orthogonal quadratic |
| **β** | (1, –2, 1) | (1, –2, 1) |
| **α** | (–0.357, 1.000) | (–0.916, 1.000) |
| **τ** | (–0.916, 0.500, –0.500) | (–0.267, 0.569, 0.300) |
| Dropout | MAR | None |

Both designs used the same mean structure. Outcomes were generated from the Gaussian location-scale model in (16)-(17), with a trajectory transition point at *t*\* = 2. Specifically, let (*k* – *t*\*)_+_ = max(*k* – *t*\*, 0) denote the post-change-point effect. For subject *i* = 1, …, *N*, we generated a binary covariate *u*_*i*_ ~ Bernoulli(0.5) and a subject-specific random scale effect 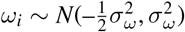 with *σ*_*ω*_ = 0.2 (see Remark 3).

For Design A, the between-subject and within-subject error variances were specified as

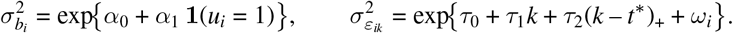

Random intercepts and within-subject errors were independently generated as 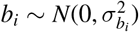 and 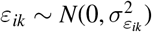, yielding

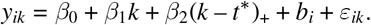

To assess the performance under missing-at-random (MAR) dropout in Design A, the observation indicator *r*_*it*_ was generated sequentially for *t* = 2, …, *K*. Conditional on being observed at visit *t* – 1, the probability of remaining observed at visit *t* followed a logistic model depending on the previously observed outcome:

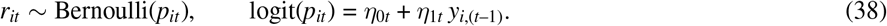

The dropout parameters were set to *η*_0*t*_ = 2.442 and *η*_1*t*_ = –0.100; the resulting missingness rates are given with **Table 2**, and dropout-parameter performance in **Supplemental Table S2**. Marginal and pairwise observation probabilities were formed as in Section 4.3.

**TABLE 2.**
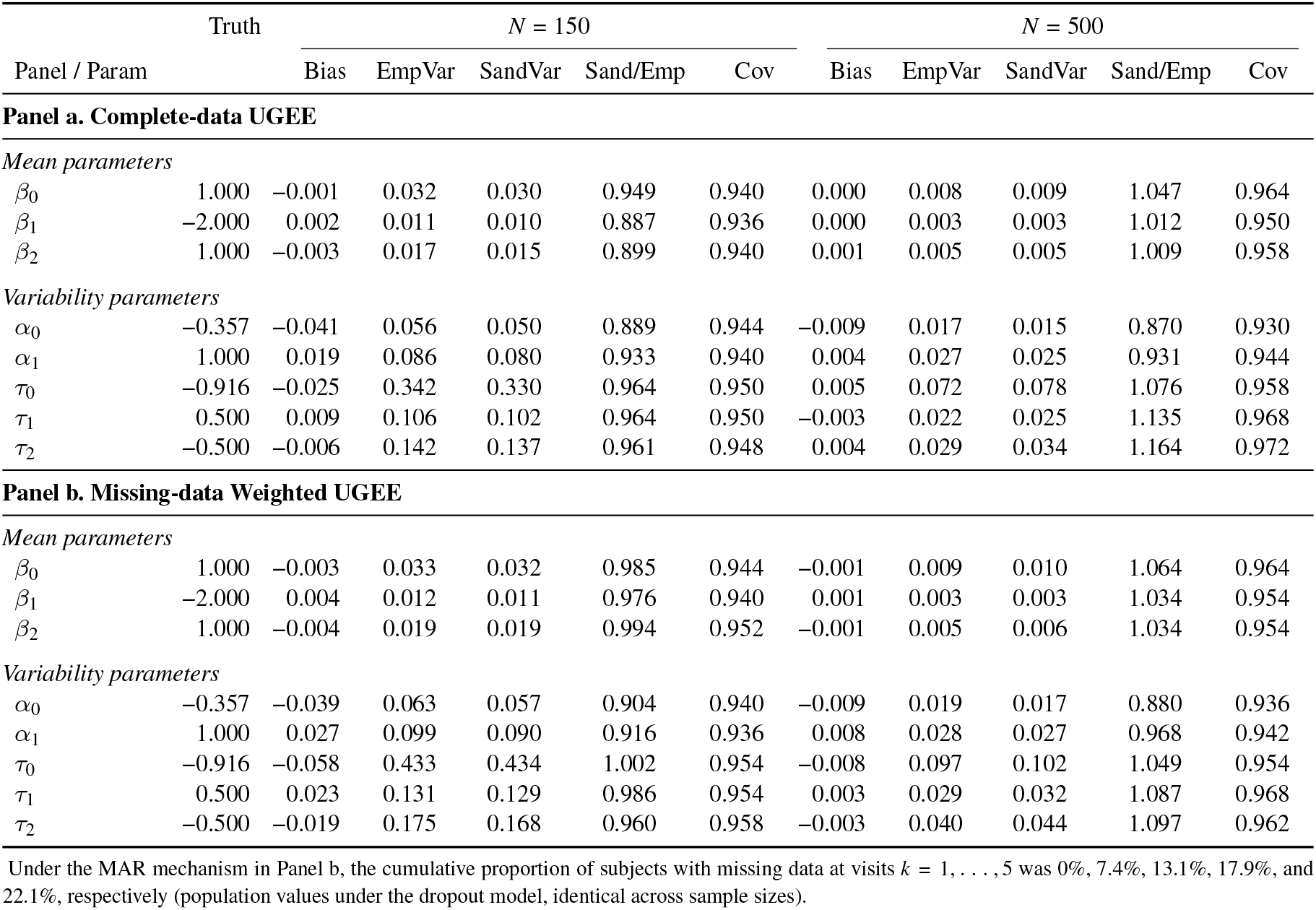
Finite-sample performance of complete-data UGEE and weighted UGEE (WUGEE) for estimation of Gaussian location-scale model parameters based on 500 Monte Carlo replications. Panel b uses pair-level weights constructed from subject-level MAR dropout. EmpVar denotes the empirical variance, SandVar the average sandwich variance estimate, Sand/Emp the ratio of average sandwich variance to empirical variance, and Cov the empirical coverage of nominal 95% Wald confidence intervals.

|  | Truth | *N* = 150 |  |  |  |  | *N* = 500 |  |  |  |  |
|---|---|---|---|---|---|---|---|---|---|---|---|
| Panel / Param |  | Bias | EmpVar | SandVar | Sand/Emp | Cov | Bias | EmpVar | SandVar | Sand/Emp | Cov |
| **Panel a. Complete-data UGEE** |  |  |  |  |  |  |  |  |  |  |  |
| *Mean parameters* |  |  |  |  |  |  |  |  |  |  |  |
| β<sub>0</sub> | 1.000 | –0.001 | 0.032 | 0.030 | 0.949 | 0.940 | 0.000 | 0.008 | 0.009 | 1.047 | 0.964 |
| β<sub>1</sub> | –2.000 | 0.002 | 0.011 | 0.010 | 0.887 | 0.936 | 0.000 | 0.003 | 0.003 | 1.012 | 0.950 |
| β<sub>2</sub> | 1.000 | –0.003 | 0.017 | 0.015 | 0.899 | 0.940 | 0.001 | 0.005 | 0.005 | 1.009 | 0.958 |
| *Variability parameters* |  |  |  |  |  |  |  |  |  |  |  |
| α<sub>0</sub> | –0.357 | –0.041 | 0.056 | 0.050 | 0.889 | 0.944 | –0.009 | 0.017 | 0.015 | 0.870 | 0.930 |
| α<sub>1</sub> | 1.000 | 0.019 | 0.086 | 0.080 | 0.933 | 0.940 | 0.004 | 0.027 | 0.025 | 0.931 | 0.944 |
| τ<sub>0</sub> | –0.916 | –0.025 | 0.342 | 0.330 | 0.964 | 0.950 | 0.005 | 0.072 | 0.078 | 1.076 | 0.958 |
| τ<sub>1</sub> | 0.500 | 0.009 | 0.106 | 0.102 | 0.964 | 0.950 | –0.003 | 0.022 | 0.025 | 1.135 | 0.968 |
| τ<sub>2</sub> | –0.500 | –0.006 | 0.142 | 0.137 | 0.961 | 0.948 | 0.004 | 0.029 | 0.034 | 1.164 | 0.972 |
| **Panel b. Missing-data Weighted UGEE** |  |  |  |  |  |  |  |  |  |  |  |
| *Mean parameters* |  |  |  |  |  |  |  |  |  |  |  |
| β<sub>0</sub> | 1.000 | –0.003 | 0.033 | 0.032 | 0.985 | 0.944 | –0.001 | 0.009 | 0.010 | 1.064 | 0.964 |
| β<sub>1</sub> | –2.000 | 0.004 | 0.012 | 0.011 | 0.976 | 0.940 | 0.001 | 0.003 | 0.003 | 1.034 | 0.954 |
| β<sub>2</sub> | 1.000 | –0.004 | 0.019 | 0.019 | 0.994 | 0.952 | –0.001 | 0.005 | 0.006 | 1.034 | 0.954 |
| *Variability parameters* |  |  |  |  |  |  |  |  |  |  |  |
| α<sub>0</sub> | –0.357 | –0.039 | 0.063 | 0.057 | 0.904 | 0.940 | –0.009 | 0.019 | 0.017 | 0.880 | 0.936 |
| α<sub>1</sub> | 1.000 | 0.027 | 0.099 | 0.090 | 0.916 | 0.936 | 0.008 | 0.028 | 0.027 | 0.968 | 0.942 |
| τ<sub>0</sub> | –0.916 | –0.058 | 0.433 | 0.434 | 1.002 | 0.954 | –0.008 | 0.097 | 0.102 | 1.049 | 0.954 |
| τ<sub>1</sub> | 0.500 | 0.023 | 0.131 | 0.129 | 0.986 | 0.954 | 0.003 | 0.029 | 0.032 | 1.087 | 0.968 |
| τ<sub>2</sub> | –0.500 | –0.019 | 0.175 | 0.168 | 0.960 | 0.958 | –0.003 | 0.040 | 0.044 | 1.097 | 0.962 |

Under the MAR mechanism in Panel b, the cumulative proportion of subjects with missing data at visits *k* = 1, . . . , 5 was 0%, 7.4%, 13.1%, 17.9%, and 22.1%, respectively (population values under the dropout model, identical across sample sizes).

In Design B, we retain the same mean structure but represent the within-subject variance on an orthogonal quadratic basis obtained by Gram-Schmidt orthogonalization of {1, *k, k*^2^} over the observed visits *k* = 1, …, *K* and rescaled to unit mean square to avoid strong collinearity ^53^. The within-subject variance was generated from 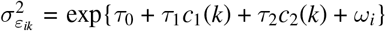, where

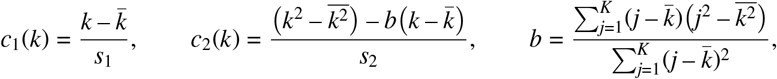

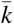 and 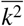 are the averages of *k* and *k*^2^ over the *K* visits, and *s*_1_, *s*_2_ > 0 are normalizing constants. The intercept, *τ*_0_, is the visit-averaged log within-subject variance, *τ*_1_ the linear trend, and *τ*_2_ the quadratic component. Design B was run under two error mechanisms: the Gaussian specification above, and a non-Gaussian mechanism with skewed random intercepts and heavy-tailed within-subject errors, specified in Section 5.4.

### 5.2 Validity of inference under the assumed model

We evaluated Monte Carlo bias, empirical variance, sandwich variance estimates, and empirical coverage of nominal 95% Wald confidence intervals. **Table 2** summarizes results for complete-data UGEE (Panel a) and weighted UGEE under MAR dropout (Panel b), where working independent correlation was used throughout.

Under complete data (Panel a), UGEE exhibited negligible bias across all parameters, with bias decreasing toward zero as sample size increased from *N* = 150 to *N* = 500. Sandwich variance estimates closely matched the empirical variance. Empirical coverage was generally near the nominal 95% level, though modest undercoverage was observed for some parameters at *N* = 150 (e.g., *β*_1_: 0.936), where sandwich variance estimates fell slightly below the empirical variance; this gap narrowed at *N* = 500 for most parameters.

Under MAR dropout (Panel b), weighted UGEE maintained comparable bias and coverage to the complete-data estimator despite moderate cumulative missingness. Sandwich variance estimates were well calibrated for both mean and variability parameters, consistent with the joint sandwich variance, which propagates uncertainty from the missingness model to inference for ***θ*** (Theorem 2). These results support reliable finite-sample performance of the proposed weighted estimating equations under the simulated MAR dropout.

### 5.3 Efficiency relative to likelihood-based inference

Under Design B we compared UGEE with the likelihood-based LSM when the Gaussian model is correctly specified, at *N* = 300 and *K ∈* {5, 10} (**Table 3**). We expect the likelihood to hold an advantage here, so this comparison isolates the efficiency price of estimating the mean and variability structures semiparametrically. We estimate relative efficiency (RE) as the LSM sampling variance divided by the UGEE sampling variance, so values below one favor the LSM. LSM coverage, reported alongside, is at or near nominal throughout, confirming that the likelihood is well calibrated when its assumptions hold.

**TABLE 3.** Efficiency of UGEE with the model-based working correlation relative to the likelihood-based Gaussian location-scale model (LSM) under correctly specified Gaussian errors, at *N* = 300. Relative efficiency (RE) is 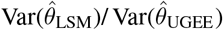; values below one favor the LSM. LSM Coverage is the empirical coverage of the LSM’s nominal 95% confidence intervals

| Param | *K* = 5 |  | *K* = 10 |  |
|---|---|---|---|---|
|  | Relative Efficiency | LSM Coverage | Relative Efficiency | LSM Coverage |
| *Mean parameters* |  |  |  |  |
| *β*<sub>0</sub> | 1.005 | 0.932 | 0.988 | 0.970 |
| *β*<sub>1</sub> | 1.000 | 0.954 | 1.000 | 0.955 |
| *β*<sub>2</sub> | 0.999 | 0.952 | 1.000 | 0.960 |
| *Variability parameters* |  |  |  |  |
| *α*<sub>0</sub> | 0.926 | 0.942 | 0.852 | 0.970 |
| *α*<sub>1</sub> | 0.987 | 0.952 | 0.966 | 0.970 |
| *τ*<sub>0</sub> | 0.541 | 0.948 | 0.327 | 0.930 |
| *τ*<sub>1</sub> | 0.569 | 0.950 | 0.444 | 0.940 |
| *τ*<sub>2</sub> | 0.586 | 0.944 | 0.487 | 0.975 |

The efficiency cost differed across the three sets of parameters. For the mean parameters, UGEE and the LSM were equally efficient (RE of 0.988 to 1.005, with no systematic change in *K*), suggesting that UGEE attains the efficiency of the correctly specified likelihood for ***β*** without imposing a distributional assumption. For the between-subject variance parameters, the efficiency loss was also modest, with RE between 0.852 and 0.987. The major efficiency tradeoff appeared for the within-subject variability parameters, where RE decreased from 0.541-0.586 at *K* = 5 to 0.327-0.487 at *K* = 10, suggesting that the correctly specified likelihood increasingly exploits distributional information for estimation of within-subject variability as the number of repeated measurements grows. It thus quantifies, in this design, the efficiency cost of not assuming a distribution.

The two approaches also differed in computational cost (**Supplemental Table S3**). At *N* = 500 and *K* = 5, UGEE required 0.331 seconds, compared with 0.669 seconds for `glmmTMB` and 585.676 seconds for `brms` under default settings (4 chains, 2000 iterations, and 1000 warmup iterations). The estimating-equation approach avoids the likelihood maximization and numerical integration required by `glmmTMB` and the MCMC sampling required by `brms`. Thus, in the examined settings, the efficiency cost of the semiparametric construction was not accompanied by a computational one.

### 5.4 Robustness to non-Gaussian errors

We further examined robustness under simultaneous violation of Gaussian assumptions and misspecification of the working correlation. In the following, random intercepts were generated from a centered and scaled *χ*^2^ distribution to induce skewness in the subject-specific effects, while within-subject errors were generated from a scaled *t* distribution with heavy tails.

Specifically, let 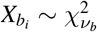 with *ν*_*b*_ = 4, and define

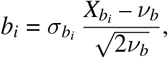

so that *b*_*i*_ has mean zero and variance 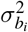. Similarly, let 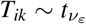 with *ν*_*ε*_ = 5, and define

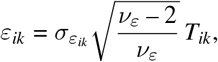

so that *ε*_*ik*_ has mean zero and variance 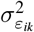 but heavier tails than Gaussian errors. Consequently, Gaussian assumptions for both random effects and residual errors were violated.

The induced within-subject correlation 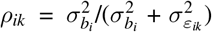 varied across time and covariate levels, so every candidate working correlation structure, applied across occasions within a pair, was misspecified. Following (26) and Section 4.1, we considered independence, exchangeable, and AR(1) structures, along with a model-based (MB) structure obtained by evaluating the pairwise covariances implied by a Gaussian location-scale model at the fitted parameter values (see **Supplement** for the derivation). Because the data were generated under non-Gaussian random effects and errors, the MB structure is also mis-specified by construction. For comparison, we also fitted the Gaussian location-scale model (LSM) via `glmmTMB`, specifying a covariate-dependent random-intercept variance and a log-linear dispersion model using the same quadratic basis. For concise-ness, we report UGEE under working independence and the MB structure alongside the LSM in the main text; results for the exchangeable and AR(1) working structures are deferred to **Supplemental Table S4**.

**Table 4** shows operating characteristic differences between UGEE and LSM. For the mean parameters, UGEE and LSM performed similarly: both remained nearly unbiased, with comparable empirical standard errors and coverage close to the nominal 0.95 level. Thus, despite substantial departures from Gaussianity, estimation and inference for the mean structure remained stable, consistent with previous findings on the robustness of classic mixed-effects models under non-Gaussian random effects ^54^.

**TABLE 4.**
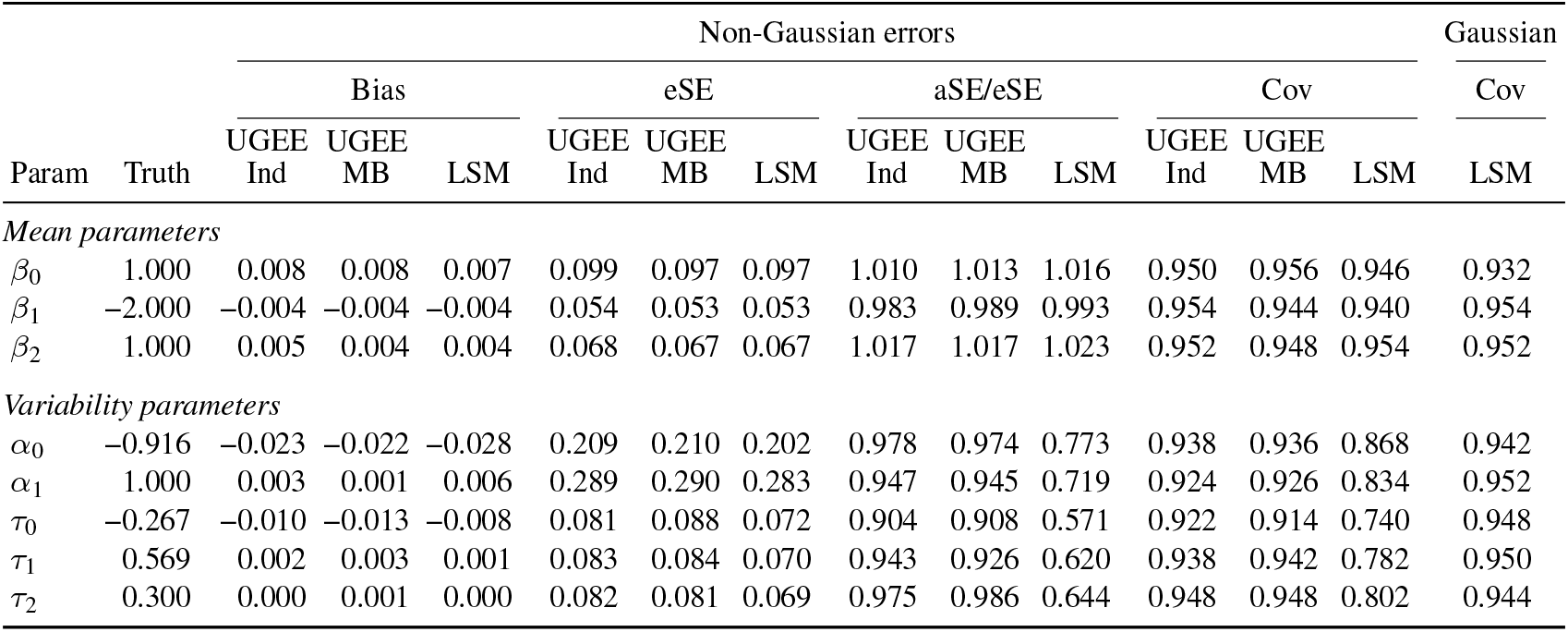
Finite-sample performance under non-Gaussian errors with *N* = 300 and *K* = 5, based on 500 Monte Carlo replications. eSE denotes the empirical standard error and aSE the average estimated standard error (sandwich-based for UGEE and model-based for LSM); aSE/eSE < 1 indicates underestimation of sampling variability. UGEE-Ind uses working independence, whereas UGEE-MB uses the model-based working covariance. The final column reports LSM coverage under the corresponding Gaussian control mechanism described in Section 5.3, reproduced from **Table 3** for comparison.

|  |  | Non-Gaussian errors |  |  |  |  |  |  |  |  |  |  |  | Gaussian |
|---|---|---|---|---|---|---|---|---|---|---|---|---|---|---|
|  |  | Bias |  |  | eSE |  |  | aSE/eSE |  |  | Cov |  |  | Cov |
| Param | Truth | UGEE<br>Ind | UGEE<br>MB | LSM | UGEE<br>Ind | UGEE<br>MB | LSM | UGEE<br>Ind | UGEE<br>MB | LSM | UGEE<br>Ind | UGEE<br>MB | LSM | LSM |
| *Mean parameters* |  |  |  |  |  |  |  |  |  |  |  |  |  |  |
| β<sub>0</sub> | 1.000 | 0.008 | 0.008 | 0.007 | 0.099 | 0.097 | 0.097 | 1.010 | 1.013 | 1.016 | 0.950 | 0.956 | 0.946 | 0.932 |
| β<sub>1</sub> | −2.000 | −0.004 | −0.004 | −0.004 | 0.054 | 0.053 | 0.053 | 0.983 | 0.989 | 0.993 | 0.954 | 0.944 | 0.940 | 0.954 |
| β<sub>2</sub> | 1.000 | 0.005 | 0.004 | 0.004 | 0.068 | 0.067 | 0.067 | 1.017 | 1.017 | 1.023 | 0.952 | 0.948 | 0.954 | 0.952 |
| *Variability parameters* |  |  |  |  |  |  |  |  |  |  |  |  |  |  |
| α<sub>0</sub> | −0.916 | −0.023 | −0.022 | −0.028 | 0.209 | 0.210 | 0.202 | 0.978 | 0.974 | 0.773 | 0.938 | 0.936 | 0.868 | 0.942 |
| α<sub>1</sub> | 1.000 | 0.003 | 0.001 | 0.006 | 0.289 | 0.290 | 0.283 | 0.947 | 0.945 | 0.719 | 0.924 | 0.926 | 0.834 | 0.952 |
| τ<sub>0</sub> | −0.267 | −0.010 | −0.013 | −0.008 | 0.081 | 0.088 | 0.072 | 0.904 | 0.908 | 0.571 | 0.922 | 0.914 | 0.740 | 0.948 |
| τ<sub>1</sub> | 0.569 | 0.002 | 0.003 | 0.001 | 0.083 | 0.084 | 0.070 | 0.943 | 0.926 | 0.620 | 0.938 | 0.942 | 0.782 | 0.950 |
| τ<sub>2</sub> | 0.300 | 0.000 | 0.001 | 0.000 | 0.082 | 0.081 | 0.069 | 0.975 | 0.986 | 0.644 | 0.948 | 0.948 | 0.802 | 0.944 |

The difference emerged for the variability parameters. Point estimation remained stable for both approaches, while LSM substantially underestimated sampling uncertainty under non-Gaussianity: its estimated vs. empirical SE ratios ranged from 0.571 to 0.773, yielding coverages of 0.740-0.868. This undercoverage was not observed under the corresponding Gaussian mechanism, where LSM coverage remained near nominal (0.932-0.954) for all parameters (**Table 3**). In contrast, UGEE sandwich standard errors more accurately matched the empirical variability (aSE vs. eSE ratio, 0.904-0.986), with coverages of 0.914-0.948. Thus, non-Gaussianity affected the LSM mainly through the standard errors of the variance parameters it models directly, which became markedly anti-conservative.

UGEE performed similarly across the four working structures: bias was negligible and coverage close to nominal for both the mean and the variability parameters, so the working structure affected precision rather than validity. For the mean model parameter estimates, precision differed little among them, as in classical identity-link GEE under a balanced and complete design ^2^. Precision of the variability model parameter estimates, obtained under a log link, did depend on the working structure (**Supplemental Table S4**), yet the independence and model-based structures were nearly identical (e.g., empirical standard error of 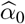 0.209 and 0.210). Working independence therefore provides a practical and computationally simple default, with no loss of precision relative to the structured alternatives in these settings.

Across the three simulation studies, semiparametric and parametric inference were nearly indistinguishable for the mean parameters: UGEE matched the correctly specified likelihood, and both remained unbiased with near-nominal coverage even when the Gaussian assumptions failed. For the variability parameters, the two approaches diverged. Under correct specification, the likelihood was more efficient, substantially so for the within-subject component and increasingly with the number of repeated measurements. Under misspecification, the same parametric structure that produced that gain was also accompanied by anti-conservative standard errors and undercoverage. Extending inference from the mean to variability therefore points to a tradeoff between precision and valid uncertainty estimates.

## 6 REAL-WORLD DATA APPLICATION

### 6.1 Background

We analyzed longitudinal systolic blood pressure (SBP) measurements from a blood pressure management program informed by the Treatment for Mild Chronic Hypertension during Pregnancy (CHAP) trial ^55,56^ at a large academic medical center among pregnant patients with chronic hypertension (*N* = 558). Each patient had up to 10 visits prior to delivery, with visit number serving as a proxy for gestational progression.

Exploratory data analysis revealed a modest decline in mean SBP that attenuated over gestation (**Figure 1(a)**). SBP variability exhibited a more pronounced nonlinear pattern, with higher variability early in pregnancy, a decline through mid-gestation, and an increase at later visits (**Figure 1(b)**). These distinct temporal patterns motivated separate characterization of the marginal mean trajectory and time-varying variability of SBP.

**FIGURE 1.**
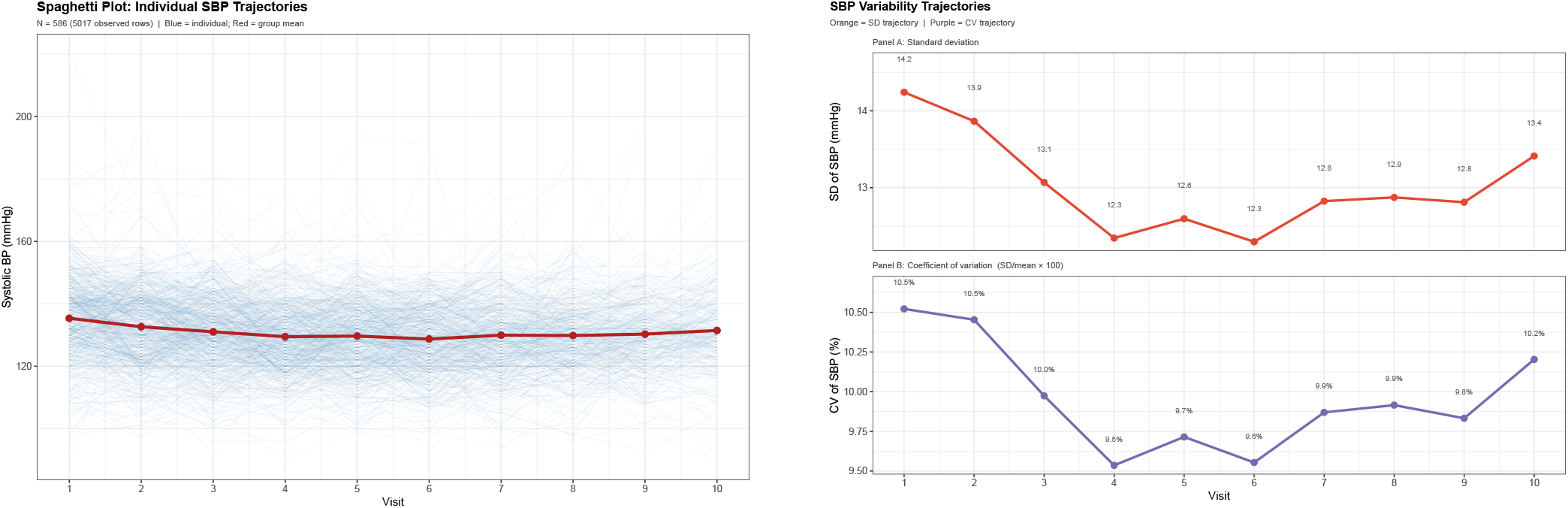
Mean and variability patterns in systolic blood pressure (SBP) during pregnancy for patients with chronic hypertension. Panel (a) shows individual SBP trajectories and the overall mean trend. Panel (b) displays the corresponding evolution of SBP variability (standard deviation and coefficient of variation) over time, illustrating substantial heterogeneity in SBP measurements across gestation (28 patients were excluded due to missing baseline covariates from the analysis cohort, with N = 558).

Our primary objective was to characterize the marginal mean, between-subject heterogeneity, and within-subject variability of SBP over gestation, while allowing the variability components to depend on clinically relevant covariates, including age, comorbid chronic kidney disease (CKD), and weight change during pregnancy.

### 6.2 Modeling Mean and Variability

We analyzed the SBP trajectories using the proposed discordance regression framework. The marginal mean model included quadratic time, baseline age, baseline weight, change from baseline weight, and CKD status:

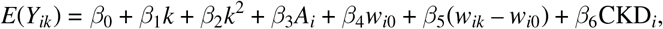

where *A*_*i*_ is baseline age, *w*_*i*0_ is baseline weight, and (*w*_*ik*_ – *w*_*i*0_) is the weight change from baseline.

To illustrate the flexibility of this framework, we used different parameterizations for the two variability components. BS heterogeneity was modeled directly through pair characteristics, whereas WS variability retained an additive individual-contribution representation analogous to the scale component of an LSM.

Specifically, the BS component was modeled using absolute covariate differences and indicators for CKD pair composition,

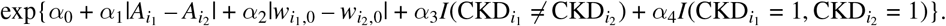

Thus, *α*_1_ and *α*_2_ characterize changes in BS heterogeneity associated with between-person differences in baseline age and weight, respectively, while *α*_3_ and *α*_4_ distinguish CKD-discordant and CKD-positive pairs from the reference group in which neither patient had CKD.

The WS component for a pair of subjects was specified as

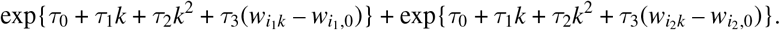

The two exponential terms are the two subjects’ own within-subject contributions, following the LSM-induced parameterization. This additive form preserves an individual-level multiplicative interpretation; for example, a one-unit increase in weight change multiplies a subject’s contribution to WS variability by exp(*τ*_3_) ^4^.

The data exhibited monotone dropout (**Supplemental Fig. S1**), which was accommodated using the weighted UGEE procedure described in Section 4.3. Under MAR, subject-specific retention probabilities were estimated from a pooled visit-to-visit logistic regression model conditioning on time, the most recently observed SBP value, CKD, baseline age, baseline weight, and the most recently observed weight. Pair-level inverse-probability weights were formed from the corresponding subject-specific retention probabilities and incorporated into the pairwise estimating equations.

### 6.3 Results

Under weighted UGEE (**Table 5**), mean SBP declined over the earlier visits 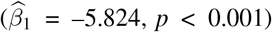, with the positive quadratic term 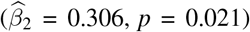 indicating that this decline attenuated over gestation. Higher baseline weight 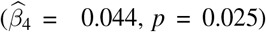 and greater weight gain from baseline 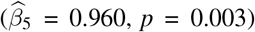 were associated with higher mean SBP, whereas baseline age and CKD showed little evidence of association.

**TABLE 5.**
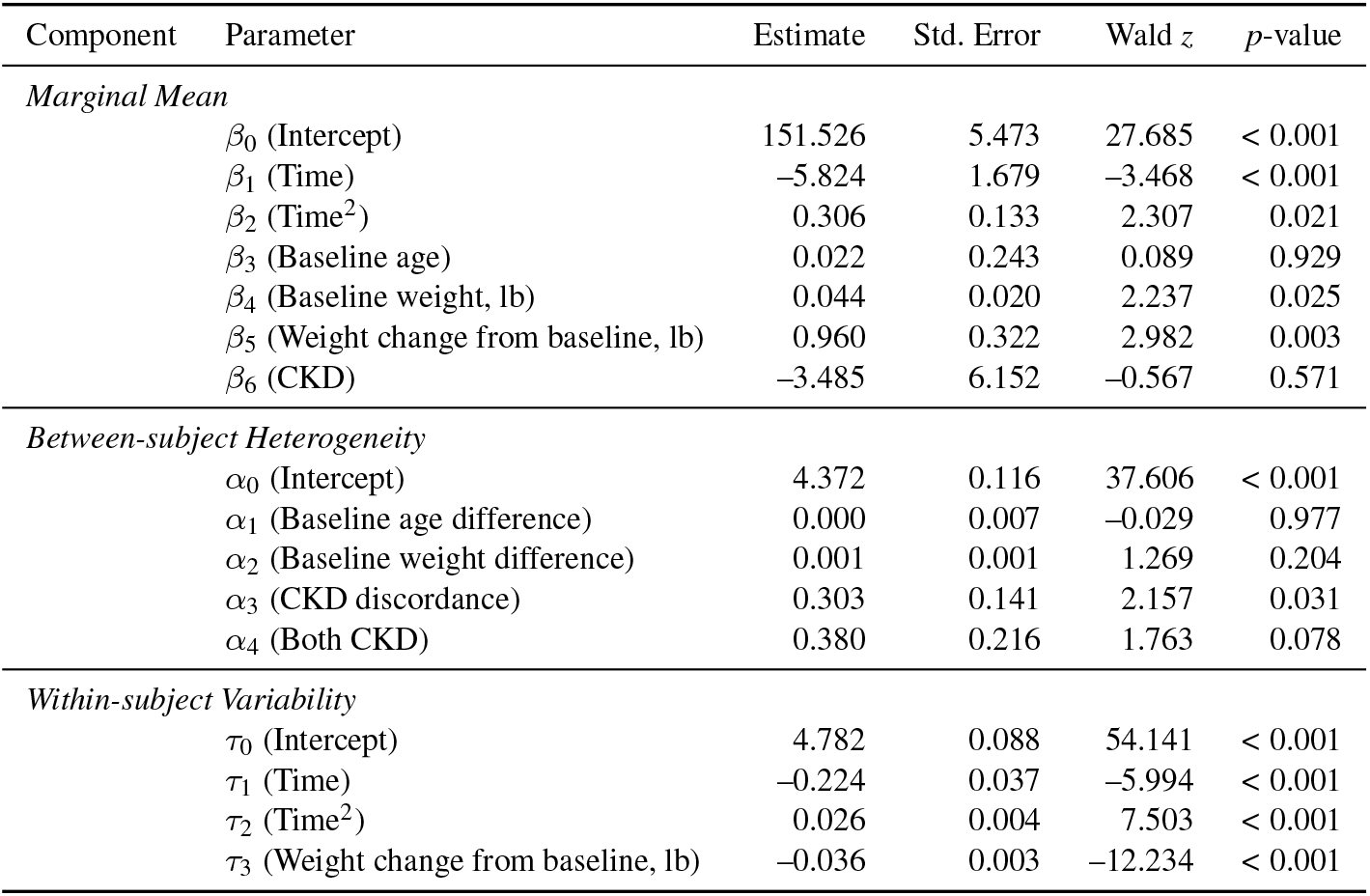
Weighted UGEE estimates for longitudinal systolic blood pressure (SBP) using inverse-probability weights from the dropout model. Parameters *β* index the marginal mean model for SBP, parameters *α* capture between-subject heterogeneity, and parameters *τ* characterize the time- and covariate-dependent within-subject variability; see Section 6.2 for the model specification.

| Component | Parameter | Estimate | Std. Error | Wald *z* | *p*-value |
|---|---|---|---|---|---|
| *Marginal Mean* |  |  |  |  |  |
|  | *β*<sub>0</sub> (Intercept) | 151.526 | 5.473 | 27.685 | < 0.001 |
|  | *β*<sub>1</sub> (Time) | –5.824 | 1.679 | –3.468 | < 0.001 |
|  | *β*<sub>2</sub> (Time<sup>2</sup>) | 0.306 | 0.133 | 2.307 | 0.021 |
|  | *β*<sub>3</sub> (Baseline age) | 0.022 | 0.243 | 0.089 | 0.929 |
|  | *β*<sub>4</sub> (Baseline weight, lb) | 0.044 | 0.020 | 2.237 | 0.025 |
|  | *β*<sub>5</sub> (Weight change from baseline, lb) | 0.960 | 0.322 | 2.982 | 0.003 |
|  | *β*<sub>6</sub> (CKD) | –3.485 | 6.152 | –0.567 | 0.571 |
| *Between-subject Heterogeneity* |  |  |  |  |  |
|  | *α*<sub>0</sub> (Intercept) | 4.372 | 0.116 | 37.606 | < 0.001 |
|  | *α*<sub>1</sub> (Baseline age difference) | 0.000 | 0.007 | –0.029 | 0.977 |
|  | *α*<sub>2</sub> (Baseline weight difference) | 0.001 | 0.001 | 1.269 | 0.204 |
|  | *α*<sub>3</sub> (CKD discordance) | 0.303 | 0.141 | 2.157 | 0.031 |
|  | *α*<sub>4</sub> (Both CKD) | 0.380 | 0.216 | 1.763 | 0.078 |
| *Within-subject Variability* |  |  |  |  |  |
|  | *τ*<sub>0</sub> (Intercept) | 4.782 | 0.088 | 54.141 | < 0.001 |
|  | *τ*<sub>1</sub> (Time) | –0.224 | 0.037 | –5.994 | < 0.001 |
|  | *τ*<sub>2</sub> (Time<sup>2</sup>) | 0.026 | 0.004 | 7.503 | < 0.001 |
|  | *τ*<sub>3</sub> (Weight change from baseline, lb) | –0.036 | 0.003 | –12.234 | < 0.001 |

For BS heterogeneity, between-person differences in baseline age and baseline weight showed little evidence of association 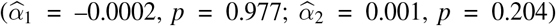. In contrast, pairs discordant on CKD status exhibited greater SBP trajectory; namely, the expected between-subject heterogeneity in SBP for a CKD-discordant pair was about 35% greater than heterogeneity than pairs in which neither patient had CKD 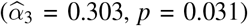, after accounting for the modeled mean for a pair in which neither patient had CKD 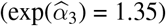. The estimate for pairs in which both patients had CKD was also positive but less precise 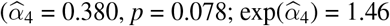, about 46% greater than the reference).

WS variability also changed nonlinearly over gestation, decreasing over the earlier visits 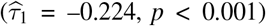 and subsequently increasing, as indicated by the positive quadratic effect 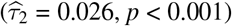. This pattern was consistent with the exploratory variability trajectory in **Figure 1(b)**. Greater weight gain was associated with lower WS variability 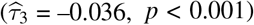, despite its positive association with mean SBP. On the multiplicative scale, each additional pound of weight gain was associated with a 3.5% lower contribution to within-subject SBP variance (exp(–0.036) = 0.965). Thus, weight change was associated differently with the level and variability of SBP, illustrating information not captured by the marginal mean trajectory alone.

As a conventional benchmark we also fit LSMs with analogous covariates under Gaussian and Student-*t* specifications. The Gaussian specification was numerically unstable; the Student-*t* specification converged, but residual diagnostics indicated departures from the fitted model, including residual heteroskedasticity and heavier-than-assumed residual tails (**Supplemental Fig. S2**). We therefore use the LSM as an empirical model-adequacy benchmark rather than for coefficient-level comparison with weighted UGEE; fitted coefficients are reported in **Supplemental Table S5**.

## 7 DISCUSSION

Joint modeling of the mean and variance in longitudinal data often encodes heterogeneity through latent random-effects variance structures, which can impose restrictions that may not align with the scientific question. We develop a semiparametric pairwise discordance regression framework that treats variability components as primary inferential targets. By modeling observable pairwise functionals directly, the proposed framework characterizes mean structure, between-subject heterogeneity, and within-subject variability without requiring a full parametric likelihood for the outcome process.

A central contribution of this work is to clarify the relationship between classical mixed-effects location-scale models and pairwise discordance regression. We show that the pairwise moment structure induced by an LSM has a particular additive form and corresponds to a restricted parameterization within the broader pairwise framework (see Section 3). For example, for a binary covariate, the LSM constrains the heterogeneous pair cell to the arithmetic mean of the two homogeneous cells, whereas the direct pair-level parameterization estimates it freely from the data. This perspective makes explicit the restrictions induced by latent variance-component formulations while allowing variability components to be parameterized according to the scientific question.

The proposed UGEE estimator provides a practical inferential tool for this pairwise framework. Because subject pairs overlap, the estimating-equation contributions are dependent, and standard independence-based asymptotic theory does not apply directly. We address this dependence using U-statistic theory and Hájek projections, yielding a sandwich variance estimator with effective sample size *N*. The resulting estimator enjoys consistency and asymptotic normality under mild regularity conditions. We further extend the framework to longitudinal missing data through inverse-probability weighted UGEE. In the setting of monotone dropout, subject-level retention models induce the weights used for the mean and pairwise variability functionals, and joint estimation of the outcome and missingness-model parameters propagates uncertainty from the weighting model into inference for ***θ***. Under the stated monotone MAR, positivity, non-interference, and correctly specified weighting-model assumptions, this construction yields valid inference for the target parameters.

The simulation studies support the finite-sample performance of the proposed estimators in the settings considered. When data were generated from the Gaussian location-scale subclass, UGEE and weighted UGEE recovered the corresponding parameters with small bias and generally well-calibrated sandwich variance estimates. Relative to the correctly specified likelihood-based LSM, UGEE showed parameter-specific efficiency tradeoffs: efficiency was nearly equivalent for mean parameters and only modestly lower for between-subject variability, whereas the likelihood-based LSM was more efficient for within-subject variability under Gaussian specification. UGEE also substantially reduced computational cost compared with MCMC-based procedures. Under non-Gaussian random effects and errors with misspecified working correlation structures, UGEE remained reasonably valid, whereas model-based LSM standard errors became anti-conservative for both variability components. This contrast illustrates the practical value of inference based on conditional moment restrictions rather than a fully specified outcome distribution.

The real-data application illustrates how the framework can separate mean, between-subject, and within-subject variability features in longitudinal systolic blood pressure (SBP) during pregnancy. In this analysis, the pair-level between-subject component allowed SBP heterogeneity to depend directly on absolute differences in baseline age and weight and on CKD pair status, while the within-subject component characterized time-varying SBP instability and its association with weight change. The finding that weight change was positively associated with mean SBP but negatively associated with within-subject SBP variability highlights the type of longitudinal feature that may be missed by analyses focused only on mean trajectories ^15^.

Several limitations and extensions merit attention. First, the current implementation uses a lag-1 discordance functional for between-subject heterogeneity, which reduces computational cost but focuses the target on adjacent-visit persistence, analogous to the successive differences underlying RMSSD (root mean square of successive differences) ^57^. Extending the construction to lag-*l* discordance for *l ∈* {1, …, *K* – 1}, or to a weighted combination across lags, would allow additional cross-time information to contribute to inference and may be particularly useful under intermittent missingness ^30,58^.

Second, the current framework assumes serially independent within-subject errors. Allowing serial dependence is a natural extension. For example, under a stationary autoregressive structure with 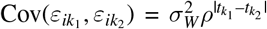, where 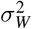 is the within-subject error variance and *ρ* the autocorrelation at unit time lag, cross-visit discordance moments would vary with lag as 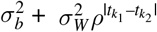, rather than reducing to a single between-subject component. Incorporating such lag-specific discordance moments into UGEE could provide a route to distinguishing stable between-subject heterogeneity, within-subject error variance 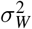, and serial correlation *ρ* in intensive longitudinal data.

Third, the current formulation uses primarily time-invariant covariates for BS heterogeneity and time-varying covariates for WS variability. This separation helps interpretation but is a modeling choice rather than a restriction of the pairwise framework. Future extensions will allow covariates to enter either or both components, accommodating predictors with both between- and within-person variation.

Fourth, as with IPW-based approaches more broadly, the missing-data framework depends on correctly specified weighting models and is less directly suited to irregularly spaced data ^31^. Augmenting weighted UGEE with an outcome-model component to obtain a doubly robust estimator ^59^ is therefore an important extension, potentially protecting against misspecification by requiring only one of the outcome or missingness models to be correctly specified. Finally, although MAR is widely adopted, future work should examine robustness through sensitivity analyses under plausible missing-not-at-random mechanisms ^60^.

More broadly, this work reframes longitudinal variability as a direct scientific target rather than a latent nuisance. This perspective is relevant in settings where both intra-individual variability and between-subject heterogeneity are of primary scientific interest, including studies of sleep ^11^, mood ^27^, heart rate ^57^, blood pressure ^28,61^, and increasingly dense data sources such as wearables and electronic health records ^12,8,13^. By defining variability targets through observable pairwise functionals, the proposed framework provides an interpretable and scalable approach for studying longitudinal heterogeneity beyond mean trajectories alone.

## Abbreviations

BS: between-subject
CHAP: Chronic Hypertension and Pregnancy
CKD: chronic kidney disease
GEE: generalized estimating equations
IPW: inverse probability weighting
LSM: location-scale model
MAR: missing at random
SBP: systolic blood pressure
UGEE: U-statistic-based generalized estimating equations
WS: within-subject.

## AUTHOR CONTRIBUTIONS

JL: Conceptualization, Methodology, Formal analysis, Software, Writing (original draft). AJ: Software, Investigation (simulation studies). SO: Resources, Data curation, Clinical interpretation, Writing (review and editing). XMT: Conceptualization, Methodology, Supervision, Writing (review and editing). JSS: Conceptualization, Supervision, Funding acquisition, Project administration, Writing (review and editing). All authors approved the final manuscript.

### Funding

This work was supported by the National Institutes of Health [R01HL094786 to Jonathan S. Schildcrout].

## DATA AVAILABILITY STATEMENT

The blood pressure data analyzed in this study contain protected health information and are available from the corresponding author upon reasonable request. Code implementing the proposed estimators, together with the scripts used to generate the reported simulation and application results, is available at https://github.com/Jinyuanliu00vumc/UGEE-location-scale.

## SUPPORTING INFORMATION

The Supplementary Material contains:

- Regularity conditions (A–F) for Theorems 1 and 2.
- Proof of Theorem 2 (consistency, asymptotic normality, and sandwich variance estimation); the proof of Theorem 1 follows standard U-statistic arguments and is omitted.
- Derivation of the Gaussian model-based working covariance for the pairwise functionals.
- Supplemental Figures S1 (monotone dropout pattern in the CHAP study) and S2 (diagnostics for the heteroskedastic Student-*t* mixed-effects model).
- Supplemental Tables S1–S5 (simulation results for *N* = 300; WUGEE dropout-parameter estimation; computation time; UGEE under four working correlation structures; and the fitted location-scale mixed model for systolic blood pressure).

