## Supplementary Material for "U-statistics-based Pairwise Discordance Regression for Joint Mean-Variability Inference in Longitudinal Studies with Missing Data"

ORIGINAL ARTICLE

**Abstract**

Longitudinal studies are central to scientific inquiry across disciplines; yet, statistical analyses have predominantly focused on mean trajectories while treating variability as a secondary or nuisance component. Increasing evidence suggests that temporal variability and heterogeneity themselves encode critical scientific information, particularly in studies with repeated measurements, heterogeneous populations, or rich covariate structures. The classical mixed-effects and location-scale model family represents variability through latent variable components but relies on strong parametric assumptions and often requires numerical approximation or computationally intensive estimation. Here we introduce the semiparametric pairwise discordance regression, a unified framework for longitudinal data that directly models pairwise contrasts to simultaneously characterize mean trajectories, between-subject heterogeneity, and within-subject temporal variability. By targeting pairwise discordances rather than latent variance components, the proposed approach enables robust inference without specifying full parametric distributions for latent components or error terms, provided the specified conditional moment restrictions hold. Pair-level covariate encodings further accommodate flexible covariate structures and allow variability parameters to be interpreted through pairwise contrasts or dissimilarities. Estimation and inference are carried out via (weighted) U-statistic-based generalized estimating equations (UGEE) with a valid sandwich variance estimator, and we extend the fitting procedure to accommodate missing-at-random dropout. We establish the asymptotic distributions of the resulting estimators under mild regularity conditions. Simulation studies demonstrate reliable finite-sample performance in the settings considered, including non-Gaussian errors and misspecified working correlation structures, whereas likelihood-based parametric fits can be sensitive to distributional assumptions or numerically unstable. An application to longitudinal blood pressure during pregnancy illustrates the framework on real data.

#### REGULARITY CONDITIONS

The following conditions are assumed throughout for the asymptotic results in Theorems 1 and 2.

##### A. Conditions on the Moment Model

**A1.** The moment restriction $E(\mathbf{f}_\mathbf{i} \mid \mathcal{F}_\mathbf{i}) = \mathbf{h}_\mathbf{i}(\boldsymbol{\theta})$ holds uniquely at $\boldsymbol{\theta}_0$; that is, $\mathbf{h}_\mathbf{i}(\boldsymbol{\theta}) = \mathbf{h}_\mathbf{i}(\boldsymbol{\theta}_0)$ almost surely implies $\boldsymbol{\theta} = \boldsymbol{\theta}_0$.
**A2.** The parameter space $\boldsymbol{\Theta} \subset \mathbb{R}^p$ is compact and convex, and $\boldsymbol{\theta}_0$ lies in its interior.

#### B. Smoothness Conditions on $\mathbf{h_i}$

**B1.** $\mathbf{h_i}(\boldsymbol{\theta})$ is continuously differentiable with respect to $\boldsymbol{\theta}$ in an open neighborhood $\mathcal{N}(\boldsymbol{\theta}_0)$ of $\boldsymbol{\theta}_0$, with Jacobian $\mathbf{G_i}(\boldsymbol{\theta}) = (\partial/\partial\boldsymbol{\theta})\mathbf{h_i}(\boldsymbol{\theta})$ continuous at $\boldsymbol{\theta}_0$.

**B2.** The Jacobian is dominated in a neighborhood of $\boldsymbol{\theta}_0$: $\sup_{\boldsymbol{\theta}\in\mathcal{N}(\boldsymbol{\theta}_0)} \|\mathbf{G_i}(\boldsymbol{\theta})\| \leq M_\mathbf{i}$ for some $M_\mathbf{i}$ satisfying $E(M_\mathbf{i}) < \infty$, ensuring that differentiation and expectation may be interchanged in the Taylor expansion of $\binom{N}{2}^{-1}\mathbf{U}_N(\boldsymbol{\theta})$.

#### C. Moment Conditions

**C1.** The residual vector has finite second moments: $E\|\mathbf{S_i}(\boldsymbol{\theta}_0)\|^2 < \infty$, where $\mathbf{S_i}(\boldsymbol{\theta}) = \mathbf{f_i} - \mathbf{h_i}(\boldsymbol{\theta})$.

**C2.** The Jacobian has finite second moments: $E\|\mathbf{G_i}(\boldsymbol{\theta}_0)\|^2 < \infty$.

**C3.** The working covariance is well-behaved: $\mathbf{V_i}$ is positive definite almost surely and $E\|\mathbf{V_i}^{-1}\| < \infty$.

#### D. Conditions for Hájek Projection

**D1.** The subject-level projected scores $\widetilde{\mathbf{v}}_{i_1}(\boldsymbol{\theta}_0) = E\{\mathbf{U}_{N,\mathbf{i}}(\boldsymbol{\theta}_0) \mid \mathcal{F}_{i_1}\}$ are independent and identically distributed across subjects $i_1 = 1, \ldots, N$, with $E\{\widetilde{\mathbf{v}}_{i_1}(\boldsymbol{\theta}_0)\} = \mathbf{0}$.

**D2.** The variance of the projected scores exists and is positive definite: $\boldsymbol{\Sigma}_v = 4\,\mathrm{Var}\{\widetilde{\mathbf{v}}_{i_1}(\boldsymbol{\theta}_0)\}$ satisfies $0 < \|\boldsymbol{\Sigma}_v\| < \infty$.

#### E. Identifiability of the Sensitivity Matrix

**E1.** The sensitivity matrix $\mathbf{B} = E\{\mathbf{G_i}\mathbf{V_i}^{-1}\mathbf{G_i}^\top\}$ is nonsingular, ensuring that the UGEE estimator $\widehat{\boldsymbol{\theta}}$ is locally unique and the asymptotic linear representation in (27) holds.

Conditions A1-A2 ensure identifiability and compactness required for uniform convergence arguments. Conditions B1-B2 justify the Taylor expansion of the UGEE and the interchange of differentiation and expectation. Conditions C1-C3 provide the moment bounds needed for the law of large numbers and central limit theorem arguments applied to the Hájek projection. Conditions D1-D2 ensure that the projected scores satisfy the classical CLT, so that the effective sample size is $N$. Condition E1 ensures invertibility of the sensitivity matrix and uniqueness of the asymptotic linear representation.

#### F. Conditions on the Missingness Mechanism

Conditions F1-F5 are additionally required for Theorem 2, which concerns the weighted estimator.

**F1.** (MAR) Conditional on being at risk, the probability of remaining observed depends only on the observed history: $\Pr(r_{ik} = 1 \mid r_{i,(k-1)} = 1, \mathbf{z}_{ik-}, y_{ik}, \ldots) = \Pr(r_{ik} = 1 \mid r_{i,(k-1)} = 1, \mathbf{z}_{ik-})$.

**F2.** (Monotonicity) $r_{i,(k+1)} = 1$ implies $r_{ik} = 1$.

**F3.** (Positivity) $p_{ik}(\boldsymbol{\eta}_0) \geq c$ almost surely for some $c > 0$ and all $i, k$.

**F4.** (Non-interference) The missingness processes are independent across subjects.

**F5.** (Correct specification) The one-step transition model holds at an interior point $\boldsymbol{\eta}_0$, and $E(\mathbf{Q_i})$ has full row rank.

Conditions F1-F4 are those stated in Section 4.3 of the main text; F1 and F5 give unbiasedness of the weighted estimating functions, F2 and F4 justify the factorizations $\pi^{(r)}_{\mathbf{i},k}$ and the at-risk restriction in (34), and F3 bounds the inverse weights.

### PROOF OF THEOREM 2

The proof of Theorem 1 follows by a multivariate extension of standard U-statistic arguments[1] and is omitted. We focus on Theorem 2, which requires additional arguments to handle the inverse probability weighted (IPW) UGEE under missing at random dropout.

We prove consistency and asymptotic normality for the joint estimator $\widehat{\vartheta} = (\widehat{\boldsymbol{\theta}}^\top, \widehat{\boldsymbol{\eta}}^\top)^\top$ defined as the solution to the stacked pairwise estimating equation (35), where the pair-level contribution is

$$\mathbf{W}_{\mathbf{i}}(\vartheta) = \begin{pmatrix} \mathbf{U}_{\mathbf{i}}^{\text{Out}}(\boldsymbol{\theta}, \boldsymbol{\eta}) \\ \mathbf{U}_{\mathbf{i}}^{\text{Mis}}(\boldsymbol{\eta}) \end{pmatrix} = \begin{pmatrix} \mathbf{G}_{\mathbf{i}} \mathbf{V}_{\mathbf{i}}^{-1} \{\mathbf{f}_{\mathbf{i}}^{\text{IPW}} - \mathbf{h}_{\mathbf{i}}(\boldsymbol{\theta})\} \\ \mathbf{Q}_{\mathbf{i}} \mathbf{M}_{\mathbf{i}}^{-1} \{\mathbf{f}_{\mathbf{i}}^{(4)} - \mathbf{q}_{\mathbf{i}}(\boldsymbol{\eta})\} \end{pmatrix}.$$

Let $\vartheta_0 = (\boldsymbol{\theta}_0^\top, \boldsymbol{\eta}_0^\top)^\top$ denote the true parameter vector.

#### Consistency

We establish unbiasedness of $\mathbf{W}_{\mathbf{i}}$ at the truth, i.e. $E\{\mathbf{W}_{\mathbf{i}}(\vartheta_0)\} = \mathbf{0}$, by treating each component separately.

*Outcome component* $\mathbf{U}_{\mathbf{i}}^{\text{Out}}$.
The IPW-weighted functional $\mathbf{f}_{\mathbf{i}}^{\text{IPW}}$ applies inverse probability weights $\pi_{\mathbf{i},k}^{-1}$ to each time-$k$ contribution (Section 4.3 of the main text). For each element indexed by visit $k \in \{1, \ldots, K\}$ and component $r \in \{1, 2, 3\}$, conditioning on the full pair-level history $\mathcal{F}_{\mathbf{i}}$ and then on the pair's outcomes $\mathbf{f}_{\mathbf{i}}$:

$$\begin{aligned} E\left( \frac{R_{\mathbf{i},k}}{\pi_{\mathbf{i},k}} \big( f_{\mathbf{i},k}^{(r)} - h_{\mathbf{i},k}^{(r)}(\vartheta_0) \big) \;\Big|\; \mathcal{F}_{\mathbf{i}} \right) &= E\left[ \big( f_{\mathbf{i},k}^{(r)} - h_{\mathbf{i},k}^{(r)} \big) \frac{E(R_{\mathbf{i},k} \mid \mathcal{F}_{\mathbf{i},k^-}, \mathbf{f}_{\mathbf{i},k^-})}{\pi_{\mathbf{i},k}} \;\Big|\; \mathcal{F}_{\mathbf{i}} \right] \qquad \text{(MAR)} \\ &= E\left[ f_{\mathbf{i},k}^{(r)} - h_{\mathbf{i},k}^{(r)}(\vartheta_0) \;\Big|\; \mathcal{F}_{\mathbf{i}} \right] = 0, \end{aligned}$$

where the MAR step uses $E(R_{\mathbf{i},k} \mid \mathcal{F}_{\mathbf{i},k^-}, \mathbf{f}_{\mathbf{i},k^-}) = \pi_{\mathbf{i},k}$, and the final equality holds by correct specification of $\mathbf{h}_{\mathbf{i}}(\boldsymbol{\theta}_0)$. Since $\mathbf{G}_{\mathbf{i}} \mathbf{V}_{\mathbf{i}}^{-1}$ is $\mathcal{F}_{\mathbf{i}}$-measurable, iterated expectations give $E\{\mathbf{U}_{\mathbf{i}}^{\text{Out}}(\vartheta_0)\} = \mathbf{0}$.

*Missingness component* $\mathbf{U}_{\mathbf{i}}^{\text{Mis}}$.
The pair-level observation indicator satisfies $R_{\mathbf{i},k} = R_{i_1,k} R_{i_2,k}$ and, under the induced pairwise model with $q_{\mathbf{i},k}(\boldsymbol{\eta}_0) = p_{i_1k}(\boldsymbol{\eta}_0) p_{i_2k}(\boldsymbol{\eta}_0)$:

$$\begin{aligned} E\Big( R_{\mathbf{i},k} - q_{\mathbf{i},k}(\boldsymbol{\eta}_0) \;\Big|\; \mathcal{F}_{\mathbf{i}} \Big) &= E\Big[ E\Big( R_{i_1,k} R_{i_2,k} \;\Big|\; R_{\mathbf{i},k-1} = 1,\, \mathcal{F}_{\mathbf{i},k^-},\, \mathbf{f}_{\mathbf{i},k^-} \Big) \;\Big|\; \mathcal{F}_{\mathbf{i}} \Big] - q_{\mathbf{i},k}(\boldsymbol{\eta}_0) \\ &= E\Big[ p_{i_1k}(\boldsymbol{\eta}_0) p_{i_2k}(\boldsymbol{\eta}_0) \;\Big|\; \mathcal{F}_{\mathbf{i}} \Big] - q_{\mathbf{i},k}(\boldsymbol{\eta}_0) = 0, \end{aligned}$$

where the second equality uses conditional independence of $R_{i_1,k}$ and $R_{i_2,k}$ given the pair history, together with correct specification of the transition probabilities. Since $\mathbf{Q}_{\mathbf{i}} \mathbf{M}_{\mathbf{i}}^{-1}$ is $\mathcal{F}_{\mathbf{i}}$-measurable, iterated expectations give $E\{\mathbf{U}_{\mathbf{i}}^{\text{Mis}}(\boldsymbol{\eta}_0)\} = \mathbf{0}$.

Combining both components yields $E\{\mathbf{W}_{\mathbf{i}}(\vartheta_0)\} = \mathbf{0}$. Consistency of $\widehat{\vartheta}$ then follows from standard arguments for U-statistic estimating equations under Conditions A–E[1,2].

#### Asymptotic Normality

Define the normalized estimating equation

$$\overline{\mathbf{W}}_N(\vartheta) := \binom{N}{2}^{-1} \sum_{\mathbf{i} \in C_2^N} \mathbf{W}_{\mathbf{i}}(\vartheta).$$

*Linearization.*
By a first-order Taylor expansion of $\overline{\mathbf{W}}_N(\widehat{\vartheta}) = \mathbf{0}$ around $\vartheta_0$, and writing $\gamma$ for the working correlation parameters:

$$\sqrt{N}(\widehat{\vartheta} - \vartheta_0) = \left(-\frac{\partial}{\partial\vartheta}\overline{\mathbf{W}}_N\right)^{-1}\left\{\sqrt{N}\,\overline{\mathbf{W}}_N(\vartheta_0) - \left(\frac{\partial}{\partial\gamma}\overline{\mathbf{W}}_N\right)^{\top}\sqrt{N}(\widehat{\gamma} - \gamma_0)\right\} + \mathbf{o}_p(1). \quad \text{(S1)}$$

*Working correlation term.*
Under Conditions A–E, $\partial\mathbf{W_i}/\partial\gamma$ is linear in the centered residuals, because the working matrices $\mathbf{V_i}$ and $\mathbf{M_i}$ enter only through $\mathbf{V_i^{-1}}$ and $\mathbf{M_i^{-1}}$ multiplying those residuals. It therefore has conditional mean zero at $(\boldsymbol{\theta}_0, \boldsymbol{\eta}_0)$, and

$$\frac{\partial}{\partial\gamma}\overline{\mathbf{W}}_N = \mathbf{o}_p(1).$$

Combined with $\sqrt{N}$-consistency of $\widehat{\gamma}$ (assumed), the working-correlation correction term in (S1) is $\mathbf{o}_p(1)$ and does not affect the limiting distribution.

*Bread matrix.*
By the law of large numbers for U-statistics (Conditions A–C),

$$\frac{\partial}{\partial\vartheta}\overline{\mathbf{W}}_N \xrightarrow{p} \widetilde{\mathbf{B}} = E\left(\frac{\partial}{\partial\vartheta}\mathbf{W_i}(\vartheta_0)\right),$$

which is block upper-triangular owing to the structure of $\mathbf{W_i}$:

$$\widetilde{\mathbf{B}} = \begin{pmatrix} \widetilde{\mathbf{B}}_{11} & \widetilde{\mathbf{B}}_{12} \\ \mathbf{0} & \widetilde{\mathbf{B}}_{22} \end{pmatrix}, \quad \text{(S2)}$$

where $\widetilde{\mathbf{B}}_{11} = -E\big(\mathbf{G_i}\mathbf{V_i^{-1}}\mathbf{G_i^{\top}}\big) = -\mathbf{B}$, $\widetilde{\mathbf{B}}_{12} = E\big(\partial\mathbf{U_i^{Out}}/\partial\boldsymbol{\eta}\big)$, and $\widetilde{\mathbf{B}}_{22} = -E\big(\mathbf{Q_i}\mathbf{M_i^{-1}}\mathbf{Q_i^{\top}}\big)$. Note that $\widetilde{\mathbf{B}}$ is defined as $E\{\partial\mathbf{W_i}/\partial(\boldsymbol{\theta}, \boldsymbol{\eta})\}$ without a leading minus, so $\widetilde{\mathbf{B}}_{11} = -\mathbf{B}$. The zero lower-left block holds because $\mathbf{U_i^{Mis}}$ does not depend on $\boldsymbol{\theta}$.

*U-statistic CLT and meat matrix.*
Define the subject-level projected score (Hájek projection):

$$\widetilde{\mathbf{v}}_{i_1}(\vartheta_0) = E\big\{\mathbf{W_i}(\vartheta_0) \mid \mathcal{F}_{i_1}\big\} = E\big\{\mathbf{W}_{(i_1, I_2)}(\vartheta_0) \mid \mathcal{F}_{i_1}\big\}, \quad \text{(S3)}$$

where $\mathcal{F}_{i_1}$ is the $\sigma$-algebra generated by subject $i_1$'s full longitudinal data and $I_2 \in \{1, \ldots, N\} \setminus \{i_1\}$ is an independent subject index. By the theory of multivariate U-statistics (Conditions A–E):

$$\sqrt{N}\,\overline{\mathbf{W}}_N(\vartheta_0) = \frac{2}{\sqrt{N}}\sum_{i_1=1}^{N}\widetilde{\mathbf{v}}_{i_1}(\vartheta_0) + \mathbf{o}_p(1) \xrightarrow{d} N(\mathbf{0},\, \boldsymbol{\Sigma}_W), \quad \boldsymbol{\Sigma}_W = 4\,\mathrm{Var}\big(\widetilde{\mathbf{v}}_{i_1}(\vartheta_0)\big).$$

Combining with (S1) and the convergence of the bread:

$$\sqrt{N}(\widehat{\vartheta} - \vartheta_0) = -\widetilde{\mathbf{B}}^{-1}\frac{2}{\sqrt{N}}\sum_{i_1=1}^{N}\widetilde{\mathbf{v}}_{i_1}(\vartheta_0) + \mathbf{o}_p(1) \xrightarrow{d} N\Big(\mathbf{0},\, \widetilde{\mathbf{B}}^{-1}\boldsymbol{\Sigma}_W\widetilde{\mathbf{B}}^{-1}\Big).$$

*Propagation of missingness uncertainty.*
Partition $\widetilde{\mathbf{B}}^{-1}$ conformably with $\vartheta = (\boldsymbol{\theta}^{\top}, \boldsymbol{\eta}^{\top})^{\top}$. By block inversion of (S2),

$$\widetilde{\mathbf{B}}^{-1} = \begin{pmatrix} \widetilde{\mathbf{B}}_{11}^{-1} & -\widetilde{\mathbf{B}}_{11}^{-1}\widetilde{\mathbf{B}}_{12}\widetilde{\mathbf{B}}_{22}^{-1} \\ \mathbf{0} & \widetilde{\mathbf{B}}_{22}^{-1} \end{pmatrix}.$$

The upper-left block of $\widetilde{\mathbf{B}}^{-1}\boldsymbol{\Sigma}_W\widetilde{\mathbf{B}}^{-1}$ is therefore

$$\begin{aligned}\boldsymbol{\Sigma}_{\boldsymbol{\theta}} &= \widetilde{\mathbf{B}}_{11}^{-1}\boldsymbol{\Sigma}_{\boldsymbol{\theta}}^{W}\widetilde{\mathbf{B}}_{11}^{-1} - \widetilde{\mathbf{B}}_{11}^{-1}\widetilde{\mathbf{B}}_{12}\widetilde{\mathbf{B}}_{22}^{-1}\boldsymbol{\Sigma}_{\boldsymbol{\eta\theta}}^{W}\widetilde{\mathbf{B}}_{11}^{-1} \\ &\quad - \widetilde{\mathbf{B}}_{11}^{-1}\boldsymbol{\Sigma}_{\boldsymbol{\theta\eta}}^{W}\widetilde{\mathbf{B}}_{22}^{-\top}\widetilde{\mathbf{B}}_{12}^{\top}\widetilde{\mathbf{B}}_{11}^{-\top} + \widetilde{\mathbf{B}}_{11}^{-1}\widetilde{\mathbf{B}}_{12}\widetilde{\mathbf{B}}_{22}^{-1}\boldsymbol{\Sigma}_{\boldsymbol{\eta}}^{W}\widetilde{\mathbf{B}}_{22}^{-\top}\widetilde{\mathbf{B}}_{12}^{\top}\widetilde{\mathbf{B}}_{11}^{-\top},\end{aligned}$$

where $\boldsymbol{\Sigma}_W = \begin{pmatrix} \boldsymbol{\Sigma}_{\boldsymbol{\theta}}^W & \boldsymbol{\Sigma}_{\boldsymbol{\theta\eta}}^W \\ \boldsymbol{\Sigma}_{\boldsymbol{\eta\theta}}^W & \boldsymbol{\Sigma}_{\boldsymbol{\eta}}^W \end{pmatrix}$. The cross-terms involving $\widetilde{\mathbf{B}}_{12}$, $\boldsymbol{\Sigma}_{\boldsymbol{\theta\eta}}^W$, and $\boldsymbol{\Sigma}_{\boldsymbol{\eta}}^W$ are generally nonzero, which readily confirms that $\boldsymbol{\Sigma}_{\boldsymbol{\theta}} \neq \widetilde{\mathbf{B}}_{11}^{-1}\boldsymbol{\Sigma}_{\boldsymbol{\theta}}^{W}\widetilde{\mathbf{B}}_{11}^{-1}$: treating $\widehat{\boldsymbol{\eta}}$ as fixed omits these terms and therefore targets $\widetilde{\mathbf{B}}_{11}^{-1}\boldsymbol{\Sigma}_{\boldsymbol{\theta}}^{W}\widetilde{\mathbf{B}}_{11}^{-1}$ rather than the asymptotic variance $\boldsymbol{\Sigma}_{\boldsymbol{\theta}}$ of $\widehat{\boldsymbol{\theta}}$.

#### Sandwich Variance Estimation

The asymptotic variance is consistently estimated by the empirical sandwich estimator. The bread is estimated by

$$\widehat{\widetilde{\mathbf{B}}} = -\binom{N}{2}^{-1} \sum_{\mathbf{i}\in C_2^N} \frac{\partial}{\partial\vartheta}\mathbf{W}_{\mathbf{i}}(\widehat{\vartheta}),$$

with blocks

$$\begin{aligned}\widehat{\widetilde{\mathbf{B}}}_{11} &= \binom{N}{2}^{-1} \sum_{\mathbf{i}\in C_2^N} \mathbf{G}_{\mathbf{i}}(\widehat{\vartheta})\,\mathbf{V}_{\mathbf{i}}^{-1}(\widehat{\gamma})\,\frac{\partial \mathbf{h}_{\mathbf{i}}}{\partial\boldsymbol{\theta}}(\widehat{\boldsymbol{\theta}}), \\ \widehat{\widetilde{\mathbf{B}}}_{22} &= \binom{N}{2}^{-1} \sum_{\mathbf{i}\in C_2^N} \mathbf{Q}_{\mathbf{i}}(\widehat{\boldsymbol{\eta}})\,\mathbf{M}_{\mathbf{i}}^{-1}(\widehat{\gamma})\,\frac{\partial \mathbf{q}_{\mathbf{i}}}{\partial\boldsymbol{\eta}}(\widehat{\boldsymbol{\eta}}).\end{aligned}$$

The Hájek-projected subject-level score is estimated by

$$\widehat{\widetilde{\mathbf{v}}}_{i_1} = \frac{1}{N-1}\sum_{i_2\neq i_1} \mathbf{W}_{(i_1,i_2)}(\widehat{\vartheta}),$$

and the meat is estimated by

$$\widehat{\boldsymbol{\Sigma}}_W = \frac{4}{N}\sum_{i_1=1}^{N} \widehat{\widetilde{\mathbf{v}}}_{i_1}\widehat{\widetilde{\mathbf{v}}}_{i_1}^{\top}.$$

The sandwich variance estimator is

$$\widehat{\boldsymbol{\Sigma}} = \widehat{\widetilde{\mathbf{B}}}^{-1}\widehat{\boldsymbol{\Sigma}}_W\widehat{\widetilde{\mathbf{B}}}^{-1},$$

and the upper-left block $\widehat{\boldsymbol{\Sigma}}_{\boldsymbol{\theta}}$ is used for inference on $\boldsymbol{\theta}$. Consistency of $\widehat{\boldsymbol{\Sigma}}$ for $\boldsymbol{\Sigma} = \widetilde{\mathbf{B}}^{-1}\boldsymbol{\Sigma}_W\widetilde{\mathbf{B}}^{-1}$ follows from consistency of $\widehat{\vartheta}$, the continuous mapping theorem, and Conditions A-E.

### MODEL-IMPLIED WORKING COVARIANCE FOR THE PAIRWISE FUNCTIONALS

Here we present the Gaussian model-based (MB) working covariance used in the simulation section of the main paper. Recall the data generating process was not Gaussian to assess the robustness of UGEE: random intercepts were drawn from a centered and scaled $\chi^2$ distribution, and within-subject errors from a scaled $t$ distribution with heavy tails. The MB specification is therefore misspecified by construction and enters the simulation as one of four working choices, alongside independence, exchangeable, and AR(1).

#### Model and notation

For subject $i$ observed at visit $k \in \{1, \ldots, K\}$, we consider a piecewise location-scale model

$$y_{ik} = \beta_0 + \beta_1 k + \beta_2(k - t^*)_+ + b_i + \varepsilon_{ik} = \boldsymbol{\beta}^{\top}\mathbf{z}_k + b_i + \varepsilon_{ik}, \tag{S4}$$

where $\mathbf{z}_k = (1, k, (k-t^*)_+)^\top$, $\boldsymbol{\beta} = (\beta_0, \beta_1, \beta_2)^\top$, and $t^*$ is a fixed knot. The errors $\varepsilon_{ik}$ are assumed independent across $i$ and across $k$, and are independent of $b_i$.

The random intercept has subject-specific variance

$$b_i \sim N(0, \sigma^2_{b,i}), \qquad \log \sigma^2_{b,i} = \alpha_0 + \alpha_1 u_i, \tag{S5}$$

where $u_i$ is a subject-level indicator and $\boldsymbol{\alpha} = (\alpha_0, \alpha_1)^\top$.

More generally, the mean and the within-subject variance need not share the same covariate sets. We write $\mathbf{z}_k$ for the mean basis above and $\mathbf{z}_k^{\text{var}} \in \mathbb{R}^3$ for the variance basis, which is the same piecewise-linear basis $\mathbf{z}_k$ under Design A by setting $\mathbf{z}_k^{\text{var}} = \mathbf{z}_k$.

In Design B, the mean structure was retained but the within-subject variance was represented on an orthogonal quadratic basis $\mathbf{z}_k^{\text{var}} = \{1, c_1(k), c_2(k)\}^\top$, obtained by Gram-Schmidt orthogonalization of $\{1, k, k^2\}$ over the observed visits $k = 1, \ldots, K$ and rescaled to unit mean square[3]; the columns are given explicitly in Section 5.1 of the main text. The within-subject variance is

$$\log \sigma^2_{\varepsilon,i}(k) = \boldsymbol{\tau}^\top \mathbf{z}_k^{\text{var}} + \omega_i, \qquad \omega_i \sim N(0, \sigma^2_\omega), \tag{S6}$$

that is, $\tau_0 + \tau_1 k + \tau_2 (k - t^*)_+$ under Design A and $\tau_0 + \tau_1 c_1(k) + \tau_2 c_2(k)$ under Design B, with $\omega_i$ assumed to be independent of $b_i$ and of $\{\varepsilon_{ik}\}$.

Because $\omega_i$ enters (S6) additively and independently of $\mathbf{z}_k^{\text{var}}$, integrating it out gives a within-subject variance that depends on $k$ alone,

$$\xi_k = E\{\sigma^2_{\varepsilon,i}(k)\} = \exp(\boldsymbol{\tau}^{*\top} \mathbf{z}_k^{\text{var}}), \qquad \tau_0^* = \tau_0 + \tfrac{1}{2}\sigma^2_\omega, \quad \tau_1^* = \tau_1, \quad \tau_2^* = \tau_2. \tag{S7}$$

The slopes are unchanged by the marginalization and the intercept is shifted by $\frac{1}{2}\sigma^2_\omega$. The conditional mean structure below depends on $(\tau_0, \sigma^2_\omega)$ only through $\tau_0^*$, so the two are not separately identified and their sum $\tau_0^*$ is estimated. In the simulations $\omega_i$ is instead drawn from $N(-\frac{1}{2}\sigma^2_\omega, \sigma^2_\omega)$, so that $E\{\exp(\omega_i)\} = 1$, hence $\tau_0^* = \tau_0$ and the reported truth for the within-subject intercept is $\tau_0$ itself.

We write $\boldsymbol{\tau} = (\tau_0^*, \tau_1, \tau_2)^\top$ throughout, and all reported intercepts are on this marginal scale. The full parameter vector is

$$\boldsymbol{\theta} = (\boldsymbol{\beta}^\top, \boldsymbol{\alpha}^\top, \boldsymbol{\tau}^\top)^\top.$$

For a subject pair $\mathbf{i} = (i_1, i_2)$, let $\mathcal{F}_\mathbf{i}$ denote the sigma-field generated by the pair covariates $(u_{i_1}, u_{i_2})$ and the observation times. All moments below are conditional on $\mathcal{F}_\mathbf{i}$ and marginal over $b_{i_1}, b_{i_2}, \omega_{i_1}, \omega_{i_2}$, and $\{\varepsilon_{ik}\}$.

#### Pairwise functionals

Write the residual $e_{ik} = y_{ik} - \boldsymbol{\beta}^\top \mathbf{z}_k$ and the pairwise residual difference

$$X_k = e_{i_1 k} - e_{i_2 k} = y_{i_1 k} - y_{i_2 k} = (b_{i_1} - b_{i_2}) + (\varepsilon_{i_1 k} - \varepsilon_{i_2 k}), \tag{S8}$$

which is free of $\boldsymbol{\beta}$. The three pairwise functionals are

$$f^{(1)}_{\mathbf{i}k} = \tfrac{1}{2}(y_{i_1 k} + y_{i_2 k}), \quad k = 1, \ldots, K; \qquad f^{(2)}_{\mathbf{i}k} = \tfrac{1}{2} X_k X_{k+1}, \quad k = 1, \ldots, K-1; \qquad f^{(3)}_{\mathbf{i}k} = \tfrac{1}{2} X_k^2, \quad k = 1, \ldots, K, \tag{S9}$$

so that stacking gives the pairwise outcome vector $\mathbf{f}_\mathbf{i} \in \mathbb{R}^{3K-1}$.

The between-subject contrast variance for the pair is

$$\sigma^2_{\Delta,\mathbf{i}} = \mathrm{Var}(b_{i_1} - b_{i_2}) = \exp(\alpha_0 + \alpha_1 u_{i_1}) + \exp(\alpha_0 + \alpha_1 u_{i_2}), \tag{S10}$$

distinct from the subject-level $\sigma^2_{b,i}$ of (S5). Since $b$ and $\varepsilon$ are assumed independent,

$$v_k \equiv \mathrm{Var}(X_k \mid \mathcal{F}_\mathbf{i}) = \sigma^2_{\Delta,\mathbf{i}} + 2\xi_k, \tag{S11}$$

and because $\varepsilon_{ik}$ is assumed serially independent, the only shared component of $X_a$ and $X_b$ at distinct times is the random intercept difference, so

$$c_{ab} \equiv \mathrm{Cov}(X_a, X_b \mid \mathcal{F}_\mathbf{i}) = \sigma^2_{\Delta,\mathbf{i}} + 2\xi_a \mathbf{1}(a = b) = \begin{cases} v_a, & a = b, \\ \sigma^2_{\Delta,\mathbf{i}}, & a \neq b. \end{cases} \tag{S12}$$

The conditional means of (S9) follow immediately:

$$h_{\mathbf{i}k}^{(1)} = \boldsymbol{\beta}^\top \mathbf{z}_k, \qquad h_{\mathbf{i}k}^{(2)} = \tfrac{1}{2} c_{k,k+1} = \tfrac{1}{2}\sigma^2_{\Delta,\mathbf{i}}, \qquad h_{\mathbf{i}k}^{(3)} = \tfrac{1}{2} c_{kk} = \tfrac{1}{2}\sigma^2_{\Delta,\mathbf{i}} + \xi_k. \tag{S13}$$

Equations (S11) through (S13) use only first and second moments, and hold regardless of the distributional form of $b_i$ and $\varepsilon_{ik}$, provided both are centered with the stated variances, $\varepsilon_{ik}$ is serially uncorrelated, and $b$ and $\varepsilon$ are independent.

#### Working covariance

*General form.*
We use the block-diagonal working covariance

$$\mathbf{V}_\mathbf{i} = \operatorname{diag}\left(\mathbf{V}_\mathbf{i}^{(1)}, \mathbf{V}_\mathbf{i}^{(2)}, \mathbf{V}_\mathbf{i}^{(3)}\right), \qquad \mathbf{V}_\mathbf{i}^{(r)} = (\mathbf{A}_\mathbf{i}^{(r)})^{1/2} \mathbf{R}_\mathbf{i}^{(r)} (\mathbf{A}_\mathbf{i}^{(r)})^{1/2}, \tag{S14}$$

where $\mathbf{A}_\mathbf{i}^{(r)}$ is diagonal with the marginal variances of the $r$th block of pairwise functionals, and $\mathbf{R}_\mathbf{i}^{(r)}$ is a working correlation matrix. Off-diagonal blocks are set to zero by construction. Note that the consistency of $\widehat{\boldsymbol{\theta}}$ requires only that the pairwise residuals have conditional mean zero at $\boldsymbol{\theta}_0$, which by (S13) is a second-moment condition, and holds for any positive definite $\mathbf{V}_\mathbf{i}$.

*Marginal variances.*
The expressions here involve fourth moments of $X_k$ and are derived under a Gaussian working model with $\sigma^2_\omega = 0$, so that $(X_1, \ldots, X_K)$ is jointly normal with covariance (S12). Isserlis' formula gives $\operatorname{Cov}(WX, YZ) = \operatorname{Cov}(W, Y)\operatorname{Cov}(X, Z) + \operatorname{Cov}(W, Z)\operatorname{Cov}(X, Y)$ for jointly centered normal $(W, X, Y, Z)$. Applying it to (S9) with (S12) yields the diagonal entries of $\mathbf{A}_\mathbf{i}^{(r)}$,

$$\operatorname{Var}(f_{\mathbf{i}k}^{(1)} \mid \mathcal{F}_\mathbf{i}) = \tfrac{1}{4} v_k = \tfrac{1}{4}\sigma^2_{\Delta,\mathbf{i}} + \tfrac{1}{2}\xi_k, \tag{S15}$$

$$\operatorname{Var}(f_{\mathbf{i}k}^{(2)} \mid \mathcal{F}_\mathbf{i}) = \tfrac{1}{4}\left(v_k v_{k+1} + \sigma^4_{\Delta,\mathbf{i}}\right), \tag{S16}$$

$$\operatorname{Var}(f_{\mathbf{i}k}^{(3)} \mid \mathcal{F}_\mathbf{i}) = \tfrac{1}{2} v_k^2. \tag{S17}$$

Note that equation (S15) involves only second moments and so does not require the Gaussian restriction, which is not the case for (S16) and (S17).

*Model-based correlations.*
The same calculation applied at distinct time indices gives the full within-block covariances. For $a, b$ in the appropriate index ranges,

$$\operatorname{Cov}(f_{\mathbf{i}a}^{(1)}, f_{\mathbf{i}b}^{(1)} \mid \mathcal{F}_\mathbf{i}) = \tfrac{1}{4} c_{ab}, \tag{S18}$$

$$\operatorname{Cov}(f_{\mathbf{i}a}^{(2)}, f_{\mathbf{i}b}^{(2)} \mid \mathcal{F}_\mathbf{i}) = \tfrac{1}{4}\left(c_{ab}\, c_{a+1,b+1} + c_{a,b+1}\, c_{a+1,b}\right), \tag{S19}$$

$$\operatorname{Cov}(f_{\mathbf{i}a}^{(3)}, f_{\mathbf{i}b}^{(3)} \mid \mathcal{F}_\mathbf{i}) = \tfrac{1}{2}\, c_{ab}^2, \tag{S20}$$

which reduce to (S15) through (S17) at $a = b$. Normalizing by the corresponding marginal variances gives the model-based working correlations

$$R_{\mathbf{i},ab}^{(1)} = \frac{c_{ab}}{\sqrt{v_a v_b}}, \qquad R_{\mathbf{i},ab}^{(3)} = \frac{c_{ab}^2}{v_a v_b}, \qquad R_{\mathbf{i},ab}^{(2)} = \frac{c_{ab}\, c_{a+1,b+1} + c_{a,b+1}\, c_{a+1,b}}{\sqrt{\left(v_a v_{a+1} + \sigma^4_{\Delta,\mathbf{i}}\right)\left(v_b v_{b+1} + \sigma^4_{\Delta,\mathbf{i}}\right)}}. \tag{S21}$$

For $a \neq b$ the first and third are $\sigma^2_{\Delta,\mathbf{i}}/\sqrt{v_a v_b}$ and $\sigma^4_{\Delta,\mathbf{i}}/(v_a v_b)$, and all three equal one at $a = b$ by construction. Each block satisfies $\Sigma_\mathbf{i}^{(r)} = (\mathbf{A}_\mathbf{i}^{(r)})^{1/2} \mathbf{R}_\mathbf{i}^{(r)} (\mathbf{A}_\mathbf{i}^{(r)})^{1/2}$, and the model-based working covariance sets $\mathbf{V}_\mathbf{i}^{(r)} = \Sigma_\mathbf{i}^{(r)}$ in (S14).

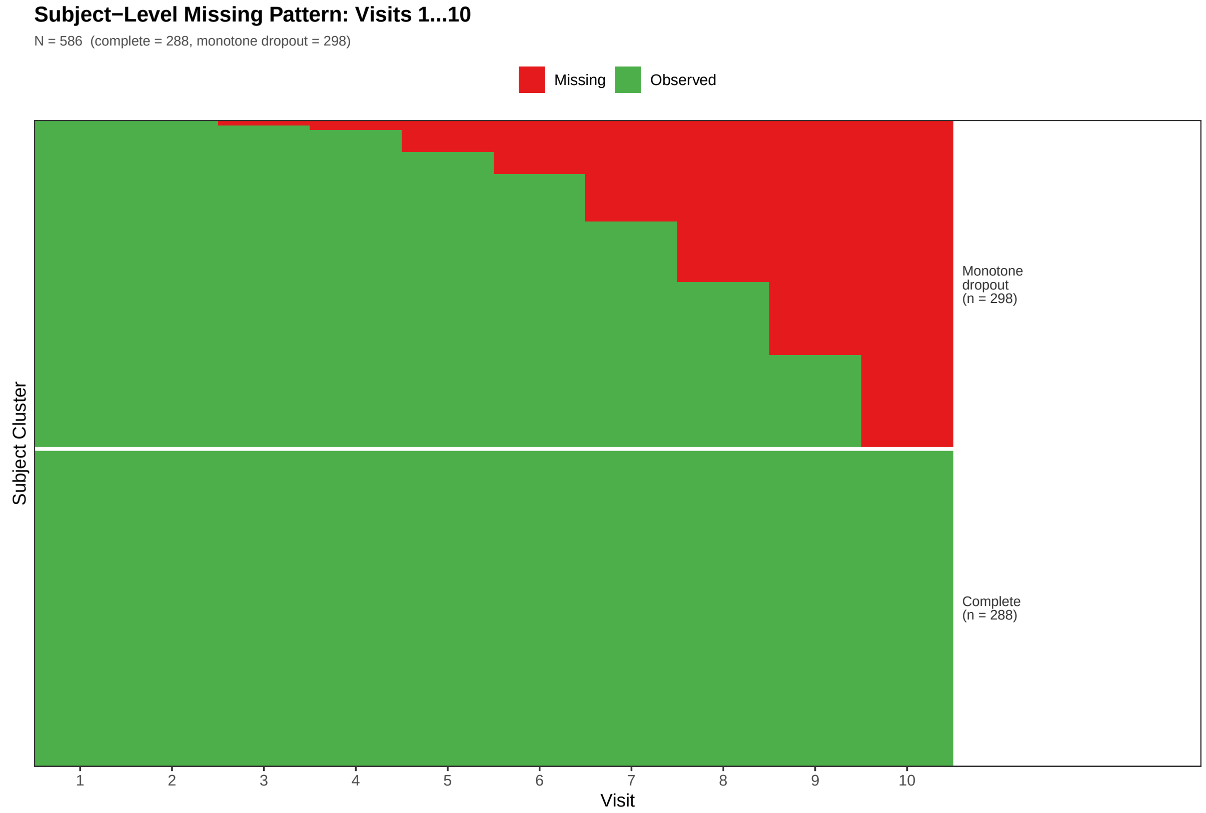


**FIGURE S1** Monotone missing data pattern of CHAP blood pressure study (28 patients were excluded due to missing baseline covariates from the analysis cohort, with $N = 558$).

### SUPPLEMENTAL FIGURES

### SUPPLEMENTAL TABLES

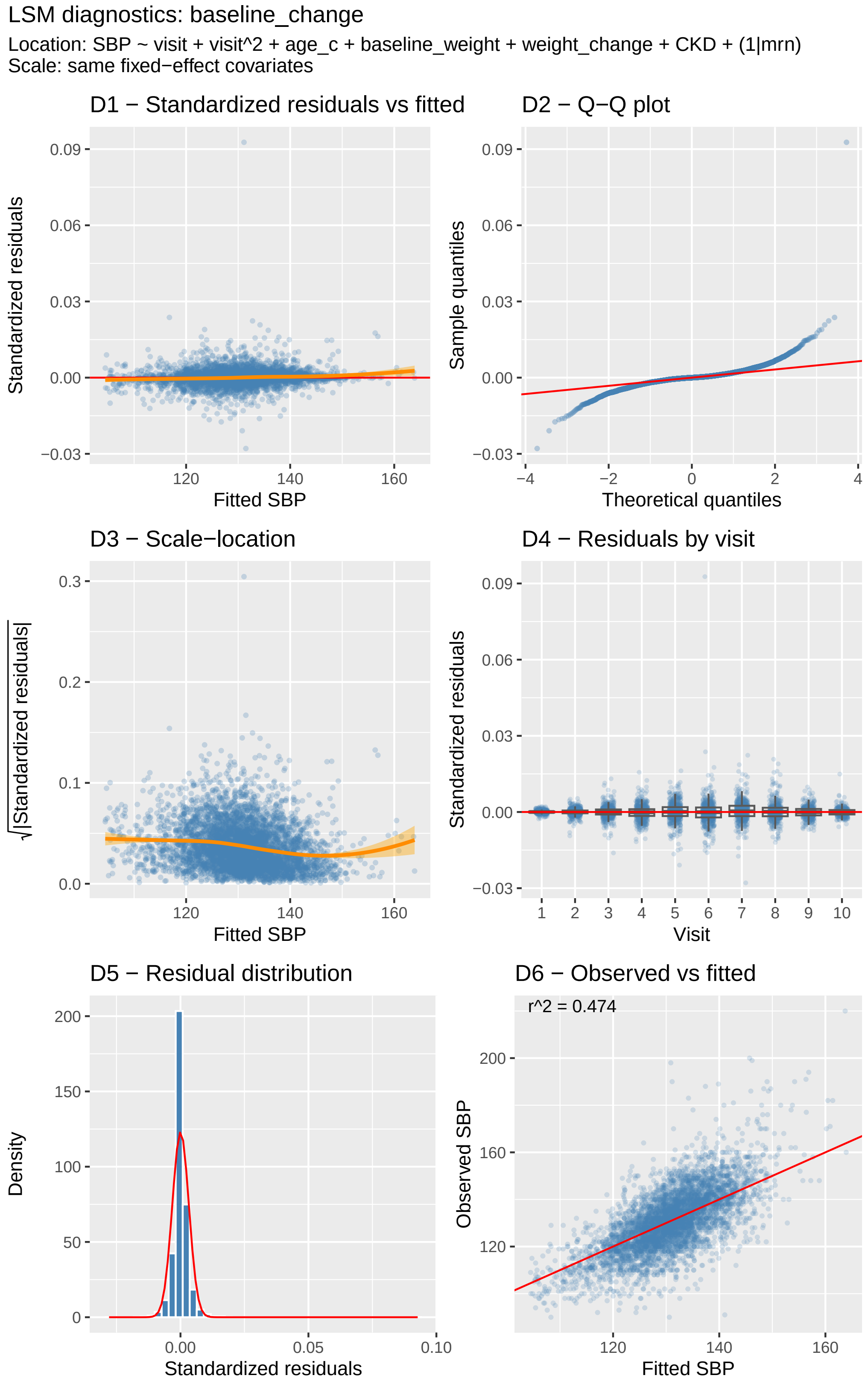


**FIGURE S2** Model diagnostics for the heteroskedastic Student-t mixed-effects model for systolic blood pressure (SBP). Panels show (D1) standardized residuals versus fitted values, (D2) QQ plot of the standardized residuals, (D3) scalelocation plot, (D4) residuals by visit number, (D5) residual distribution, and (D6) observed versus fitted SBP.

**TABLE S1** Finite-sample performance for the moderate sample size ($N = 300$) based on 500 Monte Carlo replications. EmpVar denotes the empirical variance, SandVar the average sandwich variance estimate, and Cov the empirical coverage of nominal 95% Wald confidence intervals. Under the MAR mechanism, the cumulative proportion of subjects with missing data at visits $k = 1, \ldots, 5$ was 0%, 7.4%, 13.1%, 17.9%, and 22.1%, respectively (population values under the dropout model, identical across sample sizes).

| | Complete Data | | | | Missing Data | | | |
|---|---|---|---|---|---|---|---|---|
| Param | Bias | EmpVar | SandVar | Cov | Bias | EmpVar | SandVar | Cov |
| *Mean parameters* | | | | | | | | |
| $\beta_0$ | 0.000 | 0.014 | 0.014 | 0.950 | -0.001 | 0.017 | 0.016 | 0.952 |
| $\beta_1$ | 0.000 | 0.004 | 0.005 | 0.952 | 0.005 | 0.006 | 0.006 | 0.950 |
| $\beta_2$ | 0.000 | 0.007 | 0.008 | 0.960 | -0.006 | 0.009 | 0.009 | 0.948 |
| *Variability parameters* | | | | | | | | |
| $\alpha_0$ | -0.011 | 0.025 | 0.025 | 0.948 | -0.024 | 0.029 | 0.029 | 0.952 |
| $\alpha_1$ | 0.006 | 0.045 | 0.043 | 0.944 | 0.021 | 0.047 | 0.045 | 0.940 |
| $\tau_0$ | -0.033 | 0.146 | 0.136 | 0.942 | -0.036 | 0.170 | 0.176 | 0.960 |
| $\tau_1$ | 0.018 | 0.045 | 0.043 | 0.956 | 0.016 | 0.053 | 0.054 | 0.958 |
| $\tau_2$ | -0.020 | 0.057 | 0.057 | 0.948 | -0.020 | 0.073 | 0.074 | 0.952 |
| *Dropout model parameters (WUGEE)* | | | | | | | | |
| $\eta_{0,2}$ | | – | | | 0.030 | 0.071 | 0.070 | 0.950 |
| $\eta_{1,2}$ | | – | | | -0.009 | 0.024 | 0.025 | 0.954 |
| $\eta_{0,3}$ | | – | | | 0.059 | 0.304 | 0.278 | 0.938 |
| $\eta_{1,3}$ | | – | | | -0.001 | 0.029 | 0.026 | 0.932 |
| $\eta_{0,4}$ | | – | | | 0.070 | 0.528 | 0.537 | 0.944 |
| $\eta_{1,4}$ | | – | | | 0.002 | 0.033 | 0.032 | 0.940 |
| $\eta_{0,5}$ | | – | | | 0.076 | 0.938 | 0.943 | 0.936 |
| $\eta_{1,5}$ | | – | | | 0.000 | 0.039 | 0.037 | 0.928 |

**TABLE S2** Finite-sample performance of WUGEE for estimation of dropout-model parameters under the Gaussian location-scale simulation setting with subject-level MAR dropout, based on 500 Monte Carlo replications. EmpVar denotes the empirical variance, SandVar the average sandwich variance estimate, and Cov the empirical coverage of nominal 95% Wald confidence intervals.

| | Truth | $N = 150$ | | | | $N = 500$ | | | |
|---|---|---|---|---|---|---|---|---|---|
| Parameter | | Bias | EmpVar | SandVar | Cov | Bias | EmpVar | SandVar | Cov |
| $\eta_{0,2}$ | 2.442 | 0.109 | 0.191 | 0.159 | 0.940 | 0.036 | 0.044 | 0.042 | 0.954 |
| $\eta_{1,2}$ | −0.100 | 0.008 | 0.059 | 0.052 | 0.926 | 0.006 | 0.016 | 0.015 | 0.946 |
| $\eta_{0,3}$ | 2.442 | 0.112 | 0.672 | 0.614 | 0.922 | 0.063 | 0.160 | 0.167 | 0.964 |
| $\eta_{1,3}$ | −0.100 | −0.013 | 0.075 | 0.059 | 0.884 | 0.006 | 0.015 | 0.016 | 0.950 |
| $\eta_{0,4}$ | 2.442 | 0.133 | 1.181 | 1.066 | 0.930 | 0.028 | 0.290 | 0.320 | 0.952 |
| $\eta_{1,4}$ | −0.100 | −0.008 | 0.077 | 0.061 | 0.886 | −0.002 | 0.018 | 0.019 | 0.944 |
| $\eta_{0,5}$ | 2.442 | 0.116 | 2.623 | 1.798 | 0.880 | 0.036 | 0.651 | 0.548 | 0.912 |
| $\eta_{1,5}$ | −0.100 | −0.013 | 0.099 | 0.070 | 0.860 | 0.000 | 0.024 | 0.021 | 0.920 |

Parameters $\eta_{0,t}$ and $\eta_{1,t}$ correspond to the intercept and covariate effects, respectively, in the time-specific logistic dropout models at visit $t$. Under the MAR mechanism, the cumulative proportion of subjects with missing data at visits $k = 1, \ldots, 5$ was 0%, 7.4%, 13.1%, 17.9%, and 22.1%, respectively (population values under the dropout model, identical across sample sizes).

**TABLE S3** Computation time (in seconds) for UGEE and the likelihood-based Gaussian location-scale model (LSM) across sample sizes, based on $K = 5$ repeated measurements and 500 Monte Carlo replications. LSM was fitted via two implementations: `glmmTMB` for frequentist maximum likelihood estimation and `brms` for Bayesian estimation under default settings (4 chains, 2000 iterations, 1000 warmup per chain). Times reported are averages over Monte Carlo replications.

| Sample Size | UGEE | LSM (`brms`) | LSM (`glmmTMB`) |
|---|---|---|---|
| $N = 150$ | 0.039 | 139.177 | 0.369 |
| $N = 300$ | 0.133 | 405.918 | 0.573 |
| $N = 500$ | 0.331 | 585.676 | 0.669 |

**TABLE** S4 Finite-sample performance of UGEE under four working correlation structures in the non-Gaussian heteroskedastic setting ($N = 300$, $K = 5$), based on 500 Monte Carlo replications. Columns are as in **Table 4** of the main text, with Conv the proportion of replications retained. The larger empirical standard errors under these two structures reflect a small number of heavy-tailed replications: the median-absolute-deviation-based spreads were 0.259 and 0.259 for $\widehat{\alpha}_0$ and 0.123 and 0.121 for $\widehat{\tau}_0$ (exchangeable and AR(1)), against 0.201 and 0.077 under independence.

| Param | Method | Truth | Bias | eSE | aSE/eSE | Cov | Conv |
|---|---|---|---|---|---|---|---|
| *Mean parameters* | | | | | | | |
| $\beta_0$ | UGEE-Ind | 1.000 | 0.008 | 0.099 | 1.01 | 0.950 | 1.000 |
| | UGEE-MB | | 0.008 | 0.097 | 1.01 | 0.956 | 1.000 |
| | UGEE-Exch | | 0.007 | 0.097 | 1.02 | 0.955 | 0.986 |
| | UGEE-AR1 | | 0.006 | 0.098 | 1.03 | 0.967 | 0.976 |
| $\beta_1$ | UGEE-Ind | −2.000 | −0.004 | 0.054 | 0.98 | 0.954 | 1.000 |
| | UGEE-MB | | −0.004 | 0.053 | 0.99 | 0.944 | 1.000 |
| | UGEE-Exch | | −0.004 | 0.054 | 0.99 | 0.953 | 0.986 |
| | UGEE-AR1 | | −0.003 | 0.054 | 0.99 | 0.953 | 0.976 |
| $\beta_2$ | UGEE-Ind | 1.000 | 0.005 | 0.068 | 1.02 | 0.952 | 1.000 |
| | UGEE-MB | | 0.004 | 0.067 | 1.02 | 0.948 | 1.000 |
| | UGEE-Exch | | 0.005 | 0.067 | 1.03 | 0.953 | 0.986 |
| | UGEE-AR1 | | 0.004 | 0.068 | 1.02 | 0.953 | 0.976 |
| *Variability parameters* | | | | | | | |
| $\alpha_0$ | UGEE-Ind | −0.916 | −0.023 | 0.209 | 0.98 | 0.938 | 1.000 |
| | UGEE-MB | | −0.022 | 0.210 | 0.97 | 0.936 | 1.000 |
| | UGEE-Exch | | −0.059 | 0.340 | 0.93 | 0.949 | 0.986 |
| | UGEE-AR1 | | −0.059 | 0.365 | 0.97 | 0.945 | 0.976 |
| $\alpha_1$ | UGEE-Ind | 1.000 | 0.003 | 0.289 | 0.95 | 0.924 | 1.000 |
| | UGEE-MB | | 0.001 | 0.290 | 0.95 | 0.926 | 1.000 |
| | UGEE-Exch | | 0.004 | 0.412 | 0.93 | 0.941 | 0.986 |
| | UGEE-AR1 | | −0.007 | 0.443 | 0.97 | 0.934 | 0.976 |
| $\tau_0$ | UGEE-Ind | −0.267 | −0.010 | 0.081 | 0.90 | 0.922 | 1.000 |
| | UGEE-MB | | −0.013 | 0.088 | 0.91 | 0.914 | 1.000 |
| | UGEE-Exch | | −0.008 | 0.198 | 0.75 | 0.933 | 0.986 |
| | UGEE-AR1 | | −0.004 | 0.201 | 0.78 | 0.926 | 0.976 |
| $\tau_1$ | UGEE-Ind | 0.569 | 0.002 | 0.083 | 0.94 | 0.938 | 1.000 |
| | UGEE-MB | | 0.003 | 0.084 | 0.93 | 0.942 | 1.000 |
| | UGEE-Exch | | 0.003 | 0.125 | 0.71 | 0.915 | 0.986 |
| | UGEE-AR1 | | 0.002 | 0.111 | 0.89 | 0.922 | 0.976 |
| $\tau_2$ | UGEE-Ind | 0.300 | 0.000 | 0.082 | 0.98 | 0.948 | 1.000 |
| | UGEE-MB | | 0.001 | 0.081 | 0.99 | 0.948 | 1.000 |
| | UGEE-Exch | | −0.001 | 0.092 | 0.92 | 0.957 | 0.986 |
| | UGEE-AR1 | | 0.001 | 0.092 | 0.94 | 0.953 | 0.976 |

**TABLE** S5 Estimated location-scale mixed model for systolic blood pressure (SBP) assuming a Student-$t$ distribution with identity link. The location model describes the conditional mean trajectory of SBP, while the dispersion model captures covariate effects on the residual variance. Location-model coefficients are denoted $\tilde{\beta}$ and dispersion-model coefficients $\tilde{\tau}$, to distinguish them from the pairwise-model parameters $\beta$ and $\tau$ in the main text.

| Parameter | Estimate | Std. Error | $z$ | $p$ |
|---|---|---|---|---|
| **Panel A: Location model for SBP** | | | | |
| $\tilde{\beta}_0$ (Intercept) | 137.481 | 0.645 | 213.293 | < 0.001 |
| $\tilde{\beta}_1$ (Time) | -3.023 | 0.232 | -13.009 | < 0.001 |
| $\tilde{\beta}_2$ (Time$^2$) | 0.227 | 0.020 | 11.145 | < 0.001 |
| $\tilde{\beta}_3$ (Baseline age) | -0.078 | 0.067 | -1.172 | 0.241 |
| $\tilde{\beta}_4$ (Baseline weight, lb) | 0.052 | 0.006 | 8.758 | < 0.001 |
| $\tilde{\beta}_5$ (Weight change from baseline, lb) | 0.107 | 0.019 | 5.636 | < 0.001 |
| $\tilde{\beta}_6$ (CKD) | 2.952 | 1.309 | 2.254 | 0.024 |
| **Panel B: Variance (scale) model** | | | | |
| $\tilde{\tau}_0$ (Intercept) | 2.430 | 0.048 | 50.747 | < 0.001 |
| $\tilde{\tau}_1$ (Time) | -0.125 | 0.021 | -6.011 | < 0.001 |
| $\tilde{\tau}_2$ (Time$^2$) | 0.010 | 0.002 | 5.040 | < 0.001 |
| $\tilde{\tau}_3$ (Baseline age) | 0.006 | 0.002 | 2.384 | 0.017 |
| $\tilde{\tau}_4$ (Baseline weight, lb) | 0.001 | 0.000 | 5.388 | < 0.001 |
| $\tilde{\tau}_5$ (Weight change from baseline, lb) | 0.002 | 0.001 | 1.487 | 0.137 |
| $\tilde{\tau}_6$ (CKD) | 0.127 | 0.051 | 2.509 | 0.012 |
| **Random effect** | | | | |
| $\sigma_b^2$ (random intercept variance) | 61.369 | | | |
| *Model fit statistics* | | | | |
| AIC | 38207.0 | | | |
| BIC | 38311.3 | | | |
| Log-likelihood | -19087.5 | | | |
| Observations | 5017 | | | |
| Subjects | 558 | | | |
